# NPAS4 supports drug-cue associations and relapse-like behavior through regulation of the cell type-specific activation balance in the nucleus accumbens

**DOI:** 10.1101/2022.09.04.506434

**Authors:** Brandon W. Hughes, Evgeny Tsvetkov, Benjamin M. Siemsen, Kirsten. K. Snyder, Rose Marie Akiki, Daniel J. Wood, Rachel D. Penrod, Michael D. Scofield, Stefano Berto, Makoto Taniguchi, Christopher W. Cowan

## Abstract

Use of addictive substances creates powerful drug-cue associations that often trigger relapse. Drug seeking is gated in the nucleus accumbens (NAc) by competing activation of D1 dopamine receptor-expressing medium spiny neurons (D1-MSNs) that promote, and D2 dopamine receptor-expressing neurons (D2-MSNs) that oppose, drug seeking. We show here that the ensemble of neurons in the NAc that induce the neuronal activity-regulated transcription factor, Neuronal PAS Domain Protein 4 (NPAS4), is required for cocaine-context associations. In addition, NPAS4 functions within NAc D2-MSNs to govern the activation balance of NAc D1-MSNs and D2-MSNs necessary for drug-context memories and cue-induced cocaine, but not sucrose, seeking. NPAS4 regulates drug-cue associations and preponderant D1-MSN activation by influencing a program of gene expression that blocks cocaine-induced potentiation of prefrontal cortical excitatory drive onto D2-MSNs. Together our findings reveal that NPAS4 is a key player governing NAc MSN cell-type activation balance and promoting drug-cue associations and relapse vulnerability.

## Introduction

A fundamental neurobiological mechanism underlying relapse vulnerability in substance use disorder (SUD) is the formation of powerful associations between drug-induced euphoria and the diffuse and discrete cues in the drug-taking environment ^1-3^. The nucleus accumbens (NAc) represents a key brain region that regulates motivated behavior, including relapse to drug seeking ^4,5^. In animal models of SUD, the NAc plays an essential role in mediating relapse-like behaviors, and drug experience-dependent neuronal plasticity in this region is required for future drug-seeking behavior. The NAc comprises a large neuronal population (>90%) of GABAergic long-range projecting neurons commonly referred to as medium spiny neurons (MSNs) or spiny projection neurons ^6^. MSNs are further divided into two major subpopulations that express either D1-class dopamine receptors (D1-MSNs) or D2-class dopamine receptors (D2-MSNs), which integrate motivational and reinforcement-based information from cortical and subcortical areas to facilitate goal-directed behaviors and drug seeking ^7^. In rodents, drug seeking is often studied in the drug conditioned place preference (CPP) assay or in the intravenous drug self-administration (SA) assay. CPP measures the formation of drug-context associations following experimenter-delivered (non-contingent) drug, whereas drug SA, followed by extinction training and reinstatement tests, models active drug use and relapse-like behavior. While many brain regions and circuits are involved in these complex addiction-related behaviors, the activation of glutamatergic inputs from the prelimbic PFC to the NAc core (PFC→NAc) is an essential circuit involved in cue-reinstated drug seeking ^8-13^.

Activation of NAc D1-MSNs promotes drug reward-context associations and drug-seeking behavior; whereas, activation of D2-MSNs typically opposes these behaviors ^13-23^. Indeed, optogenetic activation of D2-MSNs inhibits drug seeking by suppressing activity downstream brain regions, such as the dorsolateral ventral pallidum, through feedforward inhibition, while D1-MSN promotes cocaine seeking behaviors ^19,24-26^, and activated D2-MSNs can also directly inhibit D1-MSNs by lateral inhibition within the local microcircuit to suppress drug-related behavior ^19,27-29^. To add to the complexity of D1- and D2-MSN-mediated drug seeking, a subpopulation of neurons that induce the immediate-early gene, *Fos*, during drug conditioning are required for the expression of multiple drug behaviors ^30-32^, and the activity of these FOS+ neurons are required for future expression of many drug behaviors. While FOS is often used as a marker of recent neuronal activity, it can be induced by other stimuli, including neurotrophins and growth factors ^30,33,34^. In contrast, the intracellular calcium-dependent immediate early gene (IEG) and transcription factor, neuronal PAS domain protein 4 (NPAS4), is selectively activated by synaptic activity and L-type voltage-gated calcium channel signaling in neurons ^33,35-38^. NPAS4 in forebrain neurons maintains microcircuit homeostasis. In pyramidal neurons, it is reported to enhance inhibitory synaptic transmission, while in somatostatin-positive interneurons, it augments excitatory synaptic transmission ^33,35-37,39-42^; however, the role of NPAS4 in regulating striatal circuits and SUD-related plasticity and behavior is poorly understood. NPAS4 is induced rapidly and transiently in a small subset of NAc neurons during cocaine conditioning, and its NAc expression is required for cocaine CPP ^43^ and cocaine locomotor sensitization ^44^, but also see data herein and ^43^. Therefore, investigating the role of NAc NPAS4 in promoting drug-seeking behaviors and maintaining the D1/D2 activity balance addresses a significant knowledge gap.

## Results

### NPAS4-positive neurons during cocaine conditioning are required for future drug seeking

To test the role of the small population of NAc NPAS4-inducing neurons during cocaine conditioning on drug-context associations, we generated and validated a knock-in mouse with tamoxifen-sensitive Cre recombinase expressed with the endogenous NPAS4 coding sequence as a cleavable fusion protein under the control of the *NPAS4* gene (i.e., *NPAS4*-P2A-Cre-ERT2 or NPAS4-TRAp2a) (Figure 1A and Figure S1A-C). Following infusion of a neurotropic virus expressing a Cre-dependent, inhibitory DREADD (AAV2-DIO-hM4Di-mCherry) or negative control (AAV2-DIO-mCherry) into the NAc of the NPAS4-TRAp2a mice (Figure 1A), we administered tamoxifen (4OHT, 20 mg/kg; i.p.) immediately following either cocaine (7.5 mg/kg, i.p.) or saline conditioning sessions to induce stable expression of Gi-DREADD (or mCherry control) in the NPAS4-positive neurons (Figure 1B). Injection of clozapine-N-oxide (CNO, 5mg/kg; i.p.) in Gi-DREADD-expressing animals significantly reduced CPP (Figure 1C), without an effect in mCherry-only controls (Figure 1D). No effects were noted on locomotor activity (Figure S1D). Despite recruiting a similar number of NAc neurons (Figure S1F), Gi-DREADD-mediated inhibition of the saline-TRAPed neuronal ensemble, presumably encoding information about the saline-paired context, had no effect on cocaine CPP (Figure 1E; Figure S1E).

**Figure 1:**
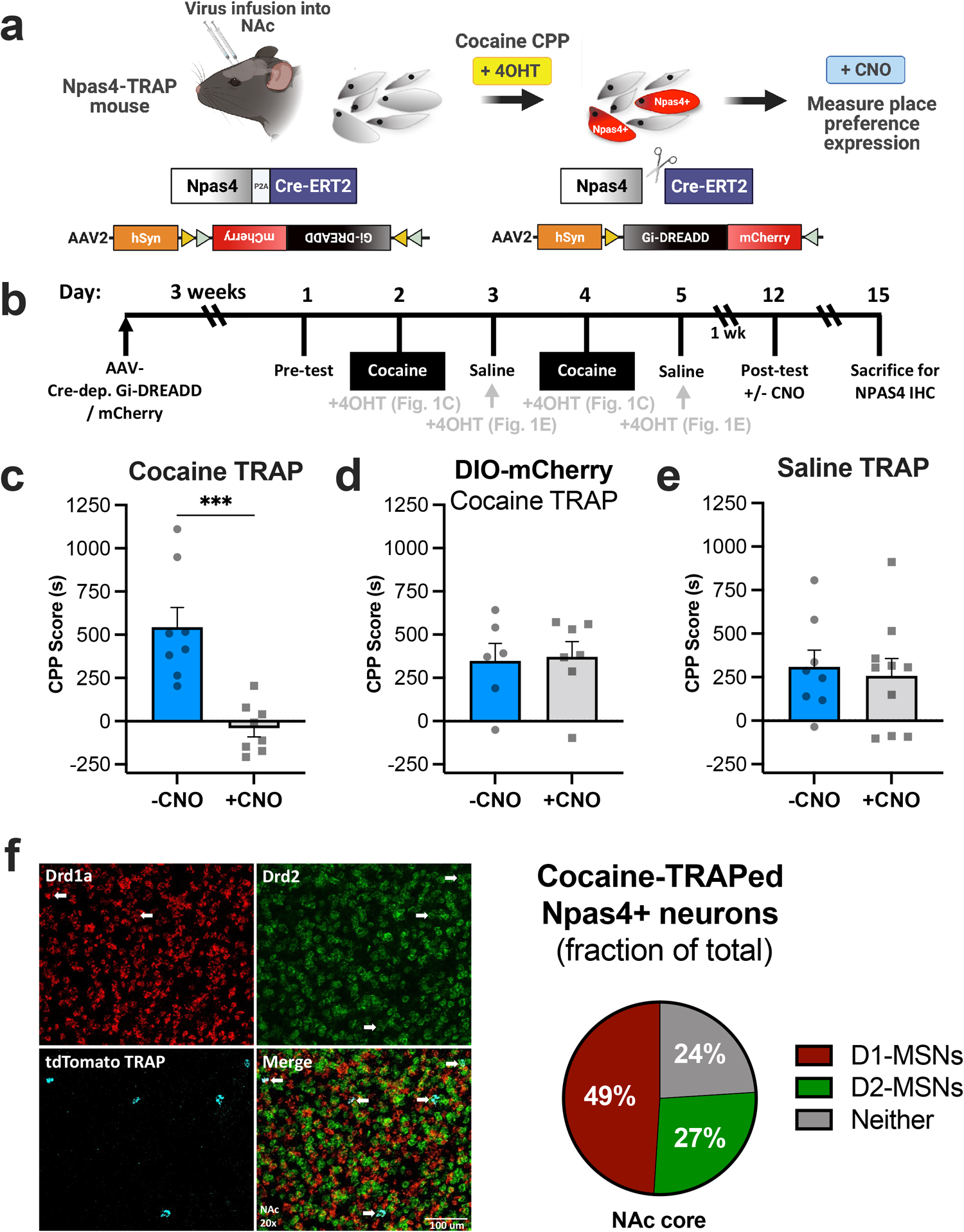
**a**, Experimental design using newly-generated NPAS4-TRAP mice. **b**, Cocaine CPP experimental timeline including 4OHT injection timing. **c**, Inhibition of the NPAS4-TRAP population captured during cocaine conditioning (n=8 mice/group) or **d**, Control virus lacking hM4Di showing no effect on cocaine CPP expression (n=6 and 7 mice/group). **e**, Inhibition of the NPAS4 ensemble activated by saline conditioning (n=8 and 10 mice/group). CPP scores calculated by time in the paired chamber – time in the unpaired chamber during the post-test (unpaired t-test; Panel C, t=4.688, df=14). **f**, Representative RNAscope images using probes for Drd1a, Drd2, and tdTomato in NAc slices of NPAS4-TRAP/Ai14 mice that received 4OHT after two cocaine conditioning sessions). **g**, Quantification of cocaine-induced NPAS4 expression (tdTomato) with D1- and D2-MSNs (n=11 mice). Data shown are mean ± SEM; ***p < 0.001.

After breeding the NPAS4-TRAp2a mice with the reporter line, *Ai14* (R26-LSL-tdTomato), and injecting the mice with tamoxifen during cocaine conditioning, we performed *in situ* hybridization with RNAscope probes that detect DA D1 or D2 receptor mRNAs. We observed that 76% of the NAc NPAS4-TRAPed neurons expressed either DA D1R (49%) or DA D2R (27%) (Figure 1F). Importantly, NPAS4-TRAp2a:Ai14 animals that did not received tamoxifen possessed one or fewer tdTomato-positive cells in the NAc (Figure S1G), confirming the high fidelity of tamoxifen-dependent recombination in the NAc. Of note, ∼40% of the NPAS4-TRAPed neurons during the 2-session cocaine or saline conditioning re-induced NPAS4 during a third cocaine conditioning session (Figure S1H), suggesting that only a subset of the NPAS4-positive neurons during initial cocaine-context conditioning is reactivated.

### NPAS4 is induced by cocaine conditioning predominantly in medium spiny neurons

D1- and D2-MSNs represent the predominant neuronal cell types in the NAc, and *in situ* hybridization (i.e., RNAscope) revealed that >75% of the NPAS4-positive neurons during cocaine conditioning express either the D1 or D2 dopamine receptors (Figure 1F). To fully characterize the NAc cell populations that express NPAS4, and to interrogate the gene expression programs regulated by both cocaine conditioning and NPAS4, we performed single nuclei RNA sequencing (snRNA-Seq) of NAc tissue isolated from mice expressing either AAV2-shNPAS4 (i.e., adeno-associated virus expressing short hairpin RNA targeting NPAS4) or AAV2-shScram virus, which does not affect NPAS4 expression. Mice were subjected to either cocaine or saline conditioning, and 24-hrs later (CPP test day), NAc tissues were rapidly extracted and processed for snRNA-seq (Figure 2A). Based on quality control stringency and unsupervised clustering, (Figure S2A-D), we defined and analyzed the NAc cell-type clusters, including subgroups of D1- and D2-MSNs, interneurons (e.g., Pvalb+, Sst+, ChAT+), and nonneuronal cells such as microglia, astrocytes, and oligodendrocytes (Figure 2B; Table S1). These clusters were defined by specific, previously published gene cluster markers of NAc cell types (Figure 2C; Figure S2D) ^45-47^, such as the enrichment of Drd1 in 3 subclusters (Figure 2C,D). NPAS4 mRNA expression was detected almost exclusively in neurons (Figure 2E), and the vast majority (∼85%) of NPAS4 was found in MSN clusters, including MSN_Drd2+, MSN_Drd1+, and MSN_Grm8+ (Figure 2D,E), the latter of which is a small, non-canonical MSN cluster associated with *Grm8* (metabotropic glutamate receptor 8) expression ^46,48-50^. We also found that, of all NAc MSN clusters, Drd2 was expressed in 27.09% of cells, and that NPAS4 mRNA was expressed in nearly 4% of all Drd2+ cells in the D2-specific clusters, compared to only 2% colocalization of NPAS4 and Drd1 in the D1-specific clusters (Interactive snRNA-seq data from this paper: https://bioinformatics-musc.shinyapps.io/Hughes_NAc_Cocaine_NPAS4/). Interestingly, cocaine conditioning produced differentially expressed genes (DEGs) in MSNs (Figure 2F), interneurons, and non-neuronal cells, including astrocytes and oligodendrocytes (Figure S2E), and that NPAS4 mRNA knockdown combined with cocaine conditioning increased the number of MSN and non-neuronal DEGs (Figure 2G), the latter presumably due to cell non-autonomous effects of neuronal NPAS4 knockdown (Figure S2E). Finally, we used RNAscope to confirm expression of NPAS4 mRNA expression in Grm8-expressing NAc MSNs (Figure S2F). Together the data suggest that NPAS4 is expressed predominantly in MSNs after saline or cocaine conditioning, and that NPAS4 influences expression of numerous genes, directly or indirectly, in multiple NAc cell populations.

**Figure 2:**
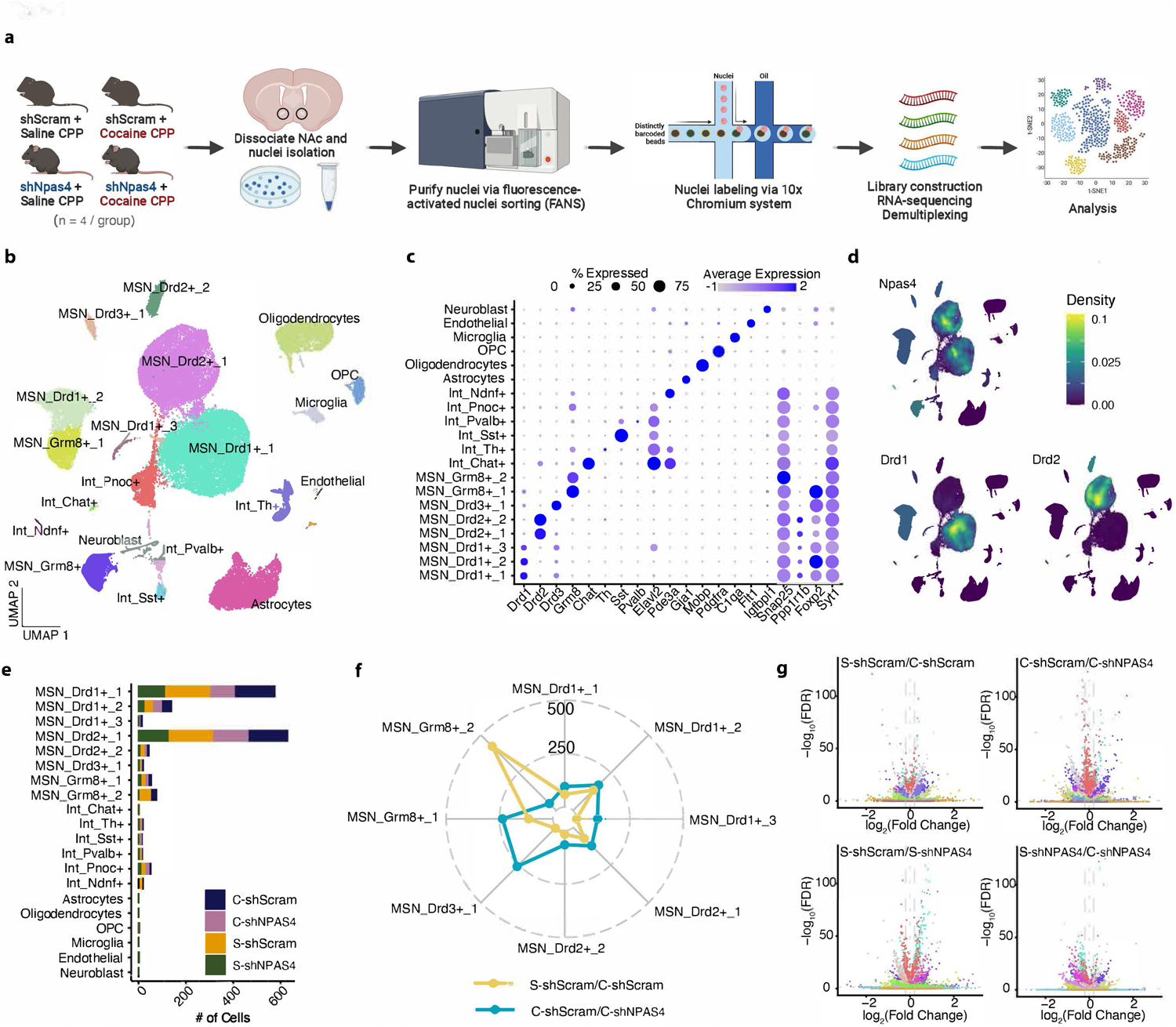
**a**, Experimental design for snRNA-seq (n=4 mice/group). **b**, Uniform manifold approximation and projection (UMAP) plot of the NAc single cells colored by cell type. Cell types were defined by known markers and confirmed by predictive modeling using a single cell NAc atlas (see methods) **c**, Dot plot showing cluster-specific markers used to generate the UMAP. Gray, low gene expression; dark blue, high gene expression. Size of circle represents the percentage of cells expressing the highlighted genes. **d**, Nebulosa plots depicting NPAS4 expression in D1-, D2-, and Grm8-MSNs. **e**, Number of NPAS4 cells in each cluster, coloured by experimental replicate. **f**, Radar plot showing the number of differentially expressed genes in each MSN cell-type identified. In yellow, the comparison between Saline and Cocaine in CPP. In blue the DEGs derived from the comparison between C-shScram vs C-shNPAS4. DEGs are defined by FDR < 0.05 and abs(log_2_(FC))>0.2. **g**, Volcano plots highlighting the transcipromic differences in each comparison, colored by cell type cluster. Y-axis corresponds to the - log_10_(FDR) whereas x-axis corresponds to the log_2_(Fold Change).

### NPAS4 functions in DA D2R-expressing neurons to promote cocaine-seeking behaviors

To test the role of NPAS4 in D1R- and D2R-expressing NAc neurons, we next generated and validated a neurotropic, Cre-dependent NPAS4 shRNA virus (AAV2-SICO-shNPAS4) (Figure 3A; Figure S3A-G) heavily modified from the first Cre-lox-regulated RNA interference system ^51^. We injected the Cre-dependent NPAS4 shRNA (shNPAS4) or a scrambled shRNA control (shScram) bilaterally in NAc of young adult D1-cre (*Drd1a*-Cre) or D2-cre (*Drd2*-Cre) male and female mice ^52,53^ (Figure 3A) and tested them in the cocaine CPP assay (Figure 3B). Selective reduction of NPAS4 in NAc D1R-expressing cells (predominantly D1-MSNs) had no significant effects on cocaine CPP (Figure 3C), cocaine-induced locomotion, or cocaine locomotor sensitization (Figure S3H). In contrast, reduction of NPAS4 in D2R-expressing cells (predominantly D2-MSNs) completely abolished cocaine CPP (Figure 3D) and showed a main effect of shNPAS4 for reduction in the locomotor sensitization assay, but a statistical trend for a miniscule reduction in locomotor sensitization to cocaine (Figure S3I), suggesting an essential D2-MSN role for NPAS4 in drug-context associations, but not in modulation of drug sensitivity.

**Figure 3:**
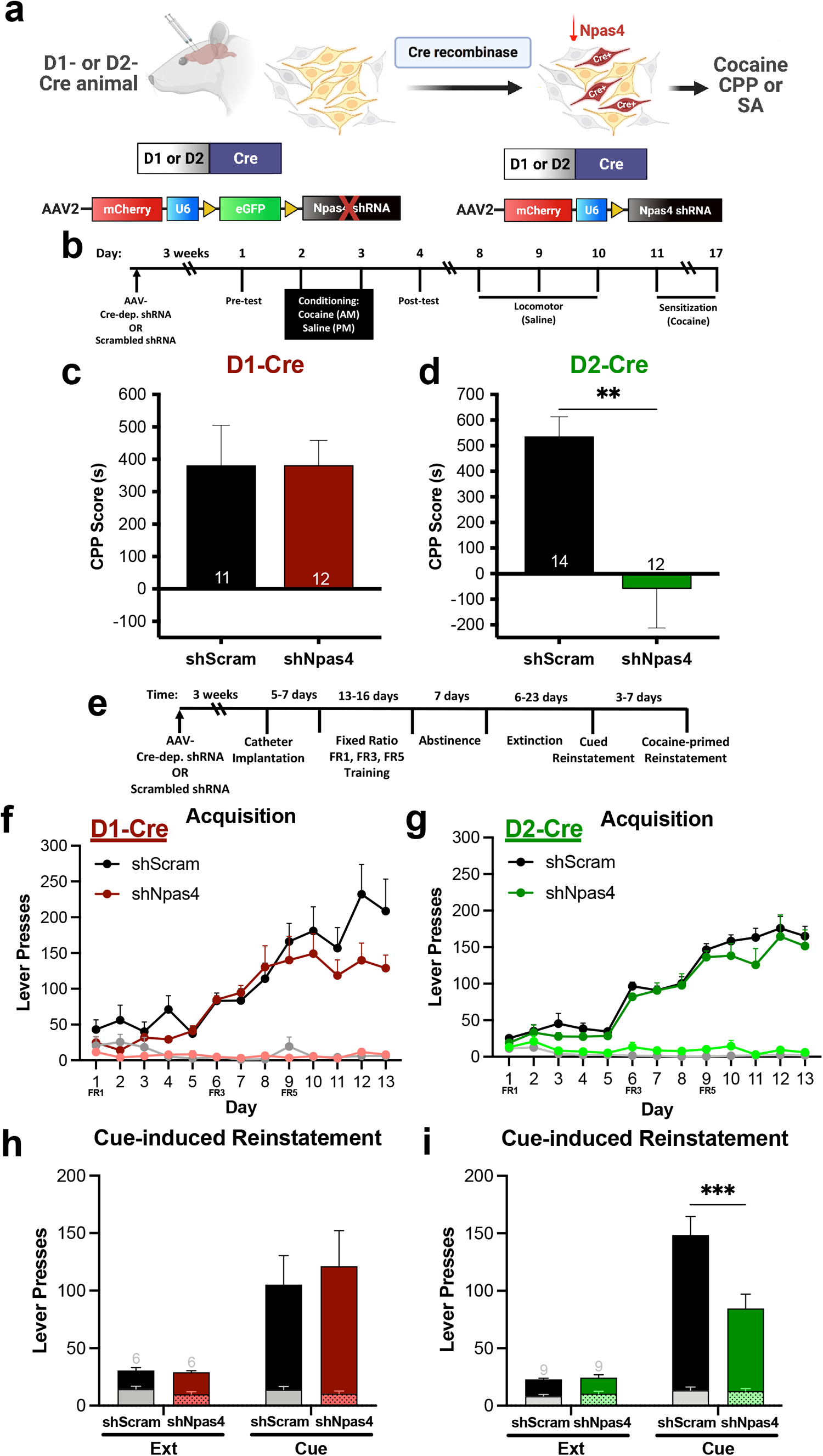
**a**, Experimental design using Cre-dependent NPAS4 shRNA in D1- or D2-Cre rats. **b**, Timeline of cocaine CPP and locomotor sensitization. **c**, Cocaine CPP with NPAS4 knockdown in NAc D1-MSNs and (D) D2-MSNs (unpaired t-test, t=3.578, df=21). **e**, Timeline of cocaine SA. **f**, Lever presses during acquisition of cocaine SA following NPAS4 knockdown in D1-Cre (n=6 rats/group) and **g**, D2-Cre rats (n=9 rats/group; 2way ANOVA with multiple comparisons; t=4.352, df=28). **h**, Cue-induced reinstatement to cocaine seeking in D1-Cre and **i**, D2-Cre rats. Data shown are mean ± SEM; **p < 0.01, ***p < 0.001.

We next employed the rat intravenous cocaine self-administration (SA) assay – a robust SUD model that incorporates volitional drug use and relapse-associated behaviors, such as cue-reinstated drug seeking ^54^. Using the pre-validated D1-Cre or D2-Cre BAC transgenic rats ^55^, we infused Cre-dependent shNPAS4 or shScram viruses into the medial NAc core (NAcore) – a NAc subregion required for cue-reinstated cocaine seeking ^4,5^. Following two weeks of daily, 2-hr cocaine SA sessions in D1- or D2-Cre rats (Figure 3E), we observed no significant differences produced by shNPAS4^NAcore^ on cocaine SA acquisition, number of drug infusions, operant discrimination of the active vs. inactive lever, or extinction of responding on the lever formerly paired with cocaine (Figure 3F,G and Figure S4A-F). We then presented the rats with the non-extinguished light and tone cues formerly linked with drug delivery and measured drug-seeking behavior (i.e., non-reinforced pressing of the drug-paired lever) (Figure 3H,I). While NAcore shNPAS4^D1-cre^ had no effect on cue-reinstated drug seeking (Figure 3H), NAcore shNPAS4^D2-Cre^ significantly reduced cue-reinstated drug seeking (Figure 3I). Of note, NAcore shNPAS4 had no effect on cocaine-primed reinstatement in either D1- or D2-Cre rats (Figure S4G,H), suggesting that NAcore NPAS4 plays a selective role in external drug-cue associations. Finally, we asked if NPAS4’s D2-MSN role in cue-reinstated seeking applied to a natural reinforcer, indicating a role in general reward associations. Using a similar operant sucrose SA design in D2-Cre rats (Figure S5A), we observed no effects of NAcore shNPAS4^D2-Cre^ on acquisition or extinction of sucrose SA (Figure S5B-E), but in contrast to reduced cue-reinstated cocaine seeking, we observed a significant increase in cue-reinstated seeking of sucrose (Figure S5F). Similar to cocaine, NAcore shNPAS4^D2-Cre^ had no effect on sucrose-primed seeking (Figure S5F). Taken together, our data reveal essential roles for NPAS4 in NAc D2-MSNs in both contingent and non-contingent models of drug seeking and opposing roles for cue-induced seeking of cocaine versus a natural reward.

### NPAS4 regulates the preponderant activation of NAc D1-MSNs following cocaine conditioning

Since activation of NAc D2-MSNs reduces drug-seeking behaviors ^7,14-21,23,24,56-59^ and loss of NPAS4 in D2-expressing neurons reduced both cocaine CPP and cue-reinstated cocaine seeking (Figures 2D,I), we next assessed the cell type-specific effects of NPAS4 knockdown on the activation balance of D1-MSNs and D2-MSNs using the activity-regulated IEG, FOS, as an indicator of recent neural activity. Following cocaine conditioning (Figure 4A) in shScram controls, we observed that ∼60% of FOS-positive NAc cells were D1-MSNs; whereas, ∼40% of FOS+ neurons were D2-MSNs (Figure 4B; 4C-D, left), as expected from prior literature ^60,61^. While NAcore shNPAS4^D1-Cre^ in NAc had no effect on the D1-MSN:D2-MSN FOS+ ratio (Figure 4C, right), NAcore shNPAS4^D2-Cre^ produced a significant shift in the D1-MSN:D2-MSN FOS+ ratio, with D2-MSNs representing the majority (39% shScram vs. 61% shNPAS4; Figure 4D, right), and a significant decrease in presumed D1-MSN FOS+ neurons (Figure 4D, right). Consistent with this observation, we also observed a significant reduction in FOS+ neurons in the dorsolateral ventral pallidum (dlVP) (Figure 4E-F), a key projection target of NAc D2-MSNs that is required for cued drug seeking ^19,21,24,26,62-64^.

**Figure 4:**
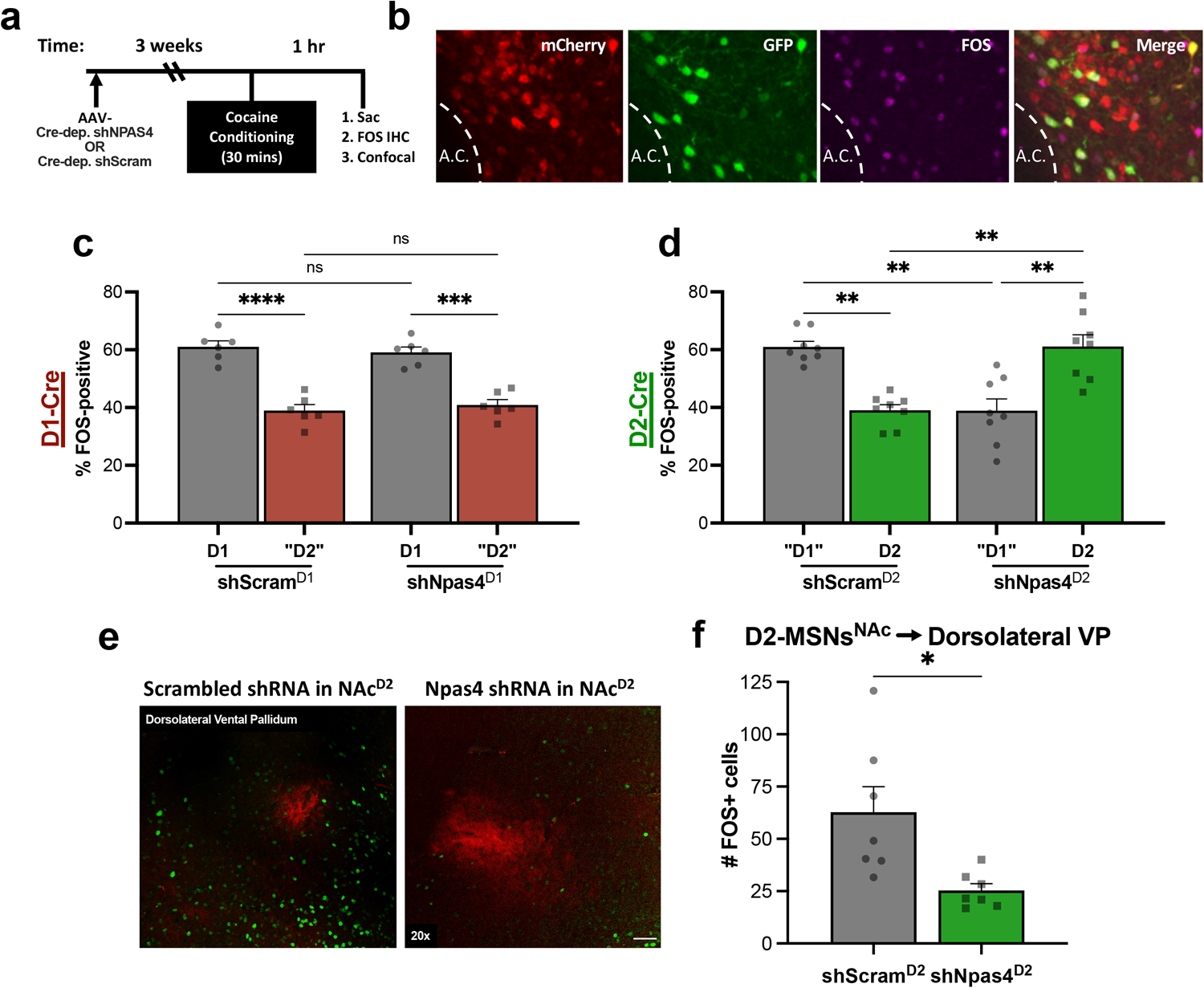
**a**, Timeline for cocaine CPP and sacrificing for FOS IHC. **b**, Representative IHC image showing Cre-dependent shRNA in the NAc of D2-Cre mice. **c**, Quantification of FOS expression in D1-(n=6 mice/group) vs. **d**, D2-Cre mice (n=8 mice/group) after cocaine CPP including effects on “putative” MSNs after cell type-specific NPAS4 knockdown (2way ANOVA with multiple comparisons, df=15). **e**, Representative image showing FOS expression in the dlVP surrounding mCherry-labeled fibers from the NAc. **f**, Quantification of FOS expression in the dlVP after NPAS4 knockdown in NAc D2-MSNs (n= 8 and 7 mice/group; unpaired t-test, t-2.973, df=12). Data shown are mean ± SEM; *p < 0.05, **p < 0.01, ***p < 0.001, ****p < 0.0001.

### NPAS4 regulates the cocaine conditioning transcriptome in NAc D2-MSNs

To determine the cell type-specific, transcriptomic influences of NAc NPAS4 on D2-MSN gene expression, we examined DEGs across all four experimental conditions (i.e., saline vs. cocaine CPP; shNPAS4 vs. shScram). All together, we detected 332 DEGs (Figure 2F; Figure 5A,D) in the D2-positive cell cluster following cocaine conditioning, and a vast majority of those DEGs were significantly upregulated. Interestingly, at this 24-hr post-conditioning timepoint, which corresponds to the transcriptional state just before CPP testing, we detected only 24 unique genes that were significantly altered in the D2-MSN cluster (Figure 5D). Analysis of the cocaine-conditioning DEGs revealed enrichment for genes linked to cocaine, amphetamine, brain development, and synapses (Figure 5B, top). In the saline-only CPP condition, we observed that shNPAS4 altered the expression of 334 genes, including *Penk, Calm1, Cartpt*, and *Camkv*, and *Cacna1h* (Figure 5A), suggesting and important role for NPAS4 on the cocaine-independent D2-MSN transcriptome. In the cocaine CPP-treated condition, shNPAS4 produced many DEGs, such as *Calm1, Cartpt, Reln, Penk*, and *Robo2* (Figure 5A; Table S2), with reported functions related to cocaine, amphetamine, synapses, neuronal projections, dendrites, and dopamine (Figure 5B, bottom). Of the DEGS regulated by both cocaine and NPAS4, specifically, there were a few genes of particular interest, including *Shisa9* and *Cartpt* (Figure 5E; top). Shisa9 is involved in regulating neuronal plasticity ^65^ while Cartpt (cocaine and amphetamine regulated transcript) overexpression negatively regulates cocaine CPP ^66^. We also interrogated genes downregulated by cocaine, but upregulated by shNPAS4, such as Penk and Tac1 (Figure 5E; bottom). Then, when we analyzed DEGs in the major D1-MSN cluster (Figure S6A), we found that shNPAS4 has a smaller impact on D1-MSNs gene expression compared to D2-MSNs (i.e., 195 in the saline condition and 96 in the cocaine condition; Figure S6D), and functional analysis of D1-MSN DEGs revealed enrichment for genes linked to cocaine, synapse organization and signaling, neuronal development, and dopamine-regulated processes (Figure S6B). Comparison of shNPAS4 DEGs in D1- and D2-MSNs revealed non-overlapping DEGs (Figure S6D), but a majority of the DEGs were observed in both cell-types, and most of the overlapping DEGs showed a similar magnitude and direction of change (Figure S6E). Finally, to assess if the shNPAS4 DEGs are potential direct targets, we compared D1- and D2-MSN DEGs with a published NPAS4 ChIP-Seq data from hippocampal neurons ^40,67^ and observed significant enrichment in promoter, intron, and distal intergenic regions (Figure S6F), with the strongest enrichment in promoters and in D2-MSNs after cocaine and shNPAS4 (Figure S6F), suggesting that some of these genes might be direct NPAS4 targets in MSNs.

**Figure 5:**
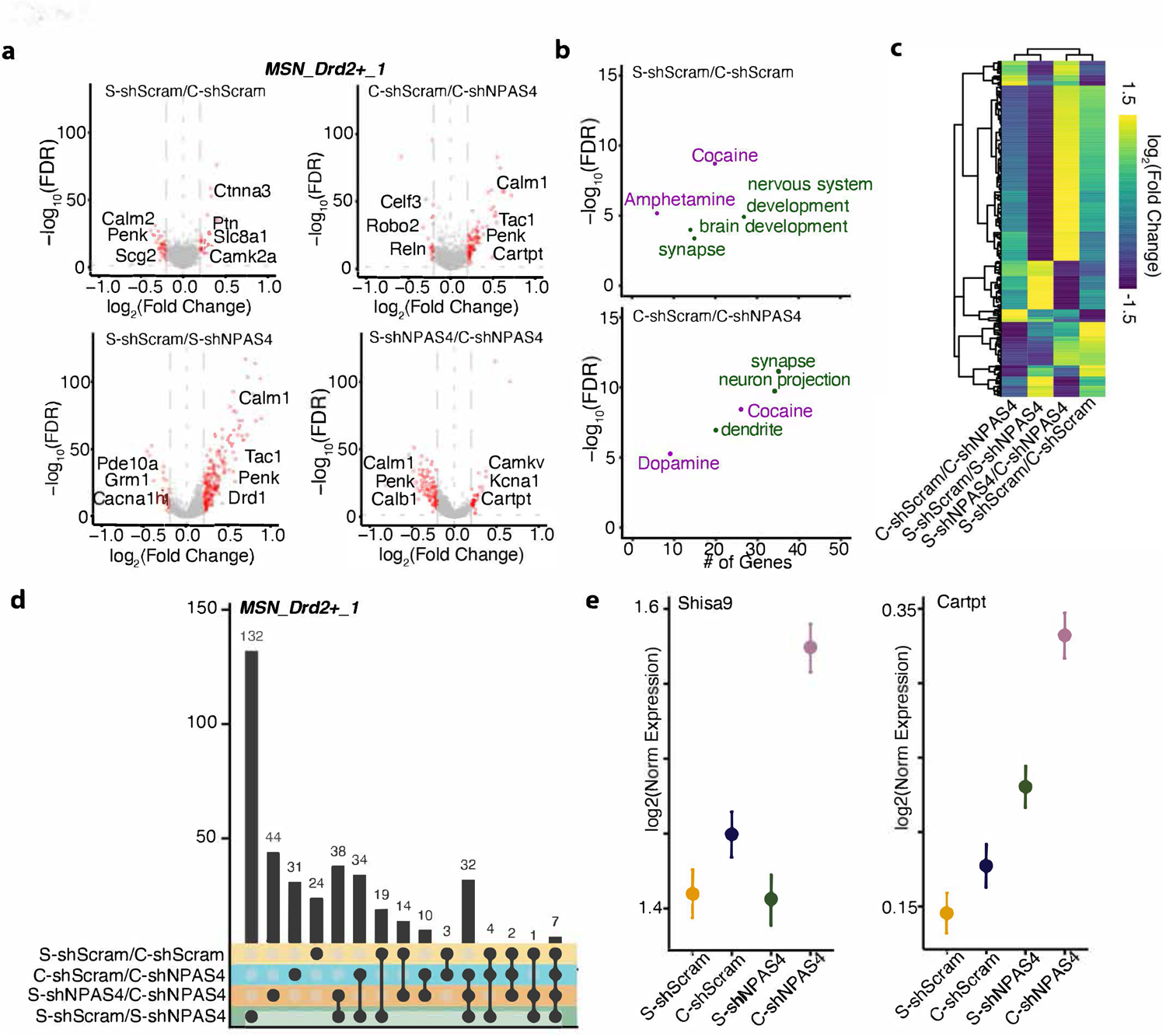
**a**, MSN_Drd2+-specific volcano plots showing genes upregulated and downregulated (DEGs) in each group comparison. Y-axis corresponds to the −log_10_(FDR) whereas x-axis corresponds to the log2(Fold Change). DEGs are defined by FDR < 0.05 and abs(log_2_(FC))>0.2. Highlighted genes of interest. **b**, Scatter plots depicting the functional enrichment for S-shScram vs C-shScram (top) and C-shScram vs C-shNPAS4 (bottom). Y-axis correspond to the −log_10_(FDR) whereas the x-axis correspond to the number of genes within the categories. In purple, categories related with drugs. In green functional categories related with molecular function. Analysis based on Fisher’s exact test. **c**, Heat map showing the fold change similarities and differences between the 4 comparisons in D2-MSNs. Hirechical clustering based on pearson’s correlation. **d**, Upset plot showing overlapping DEGs between groups in D2-MSNs. The bar graph on the top shows the number of intersections. **e**, Errorplot showing the median of the expression for DEGs in D2-MSNs significantly upregulated ONLY in the “Cocaine CPP + shNPAS4” group, Shisa9 (top left) and Cartpt (top right), as well as genes specifically downregulated by cocaine and upregulated by shNPAS4, Penk (bottom left) and Tac1 (bottom right). X-axis correspond to the experimental categories whereas the Y-axis the scaled expression level. Error bars correspond to the standard error of the mean.

### NPAS4 blocks cocaine conditioning-induced dendritic spine growth and strengthening of prelimbic cortical inputs onto NAc D2-MSNs

Since reduction of NPAS4 in NAc D2-MSNs: (1) reduces cocaine CPP and cue-reinstated cocaine seeking, (2) inverts the balance of D1:D2-MSN activation during cocaine conditioning, (3) reduces activation of essential downstream brain regions needed for drug seeking, and (4) significantly alters expression of genes linked to synapses, dendrites, neuron projections, and cocaine, we next examined NPAS4’s role in NAc D2-MSNs on glutamatergic dendritic spines – the location of most excitatory synapses on MSNs. Using a cre-dependent viral approach in D2-cre mice (Figure 6A), we found that NPAS4 knockdown in mice conditioned to cocaine, but not saline (Figure 6A), produced a significant increase in D2-MSN dendritic spine density (Figure 6B,C), and that this increase in density was observed mostly in spines with a head diameter <0.5mm (Figure S7A), suggesting that NPAS4 blocks the formation of new “thin”-type spines, which are highly motile and plastic ^68,69^. Interestingly, in saline-treated mice, no D2-MSN spine changes were observed with shNPAS4, indicating a specific interaction between NPAS4 and cocaine experience.

**Figure 6:**
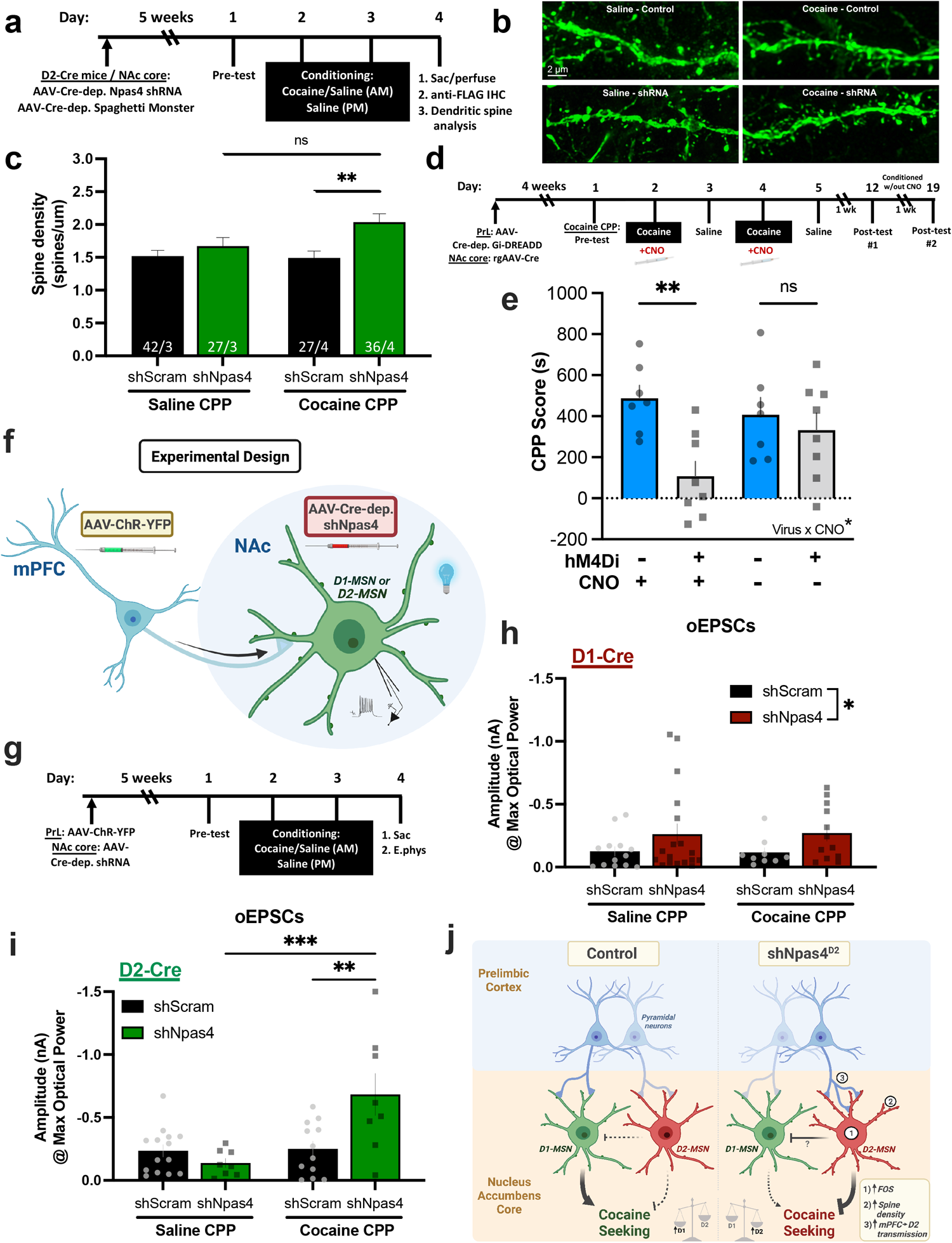
**a**, Timeline for behavior and dendritic spine labeling. **b**, Representative images of dendritic spines on NAc D2-MSNs. **c**, Quantification of spine density on D2-MSNs after cocaine (n=27 and 36 dendritic segments/group) and saline CPP (n=42 and 27 dendritic segments/group). **d**, Timeline for cocaine CPP experiment inhibiting the PrL-Nac core circuit and **e**, behavioral results following cocaine CPP (2way ANOVA with multiple comparisons). **f**, Experimental design of stimulating mPFC afferents to Nac MSNs. **g**, Timeline of cocaine CPP before electrophysiology. **h**, Evoked EPSCs onto D1-MSNs and **I**, D2-MSNs from the mPFC after NPAS4 knockdown in the corresponding cell type (2way ANOVA with multiple comparisons; D1-Cre mice: shScram n=13 and 9 cells/group, shNPAS4 n=18 and 12 cells/group; D2-Cre mice: shScram n=14 and 12 cells/group, shNPAS4 n=8 cells/group). **j**, Graphical illustration showing effects of NAc shNPAS4^D2^ on MSN activity, dendritic spines density, excitatory transmission from the mPFC, and subsequent cocaine seeking. Data shown are mean ± SEM; *p < 0.05, **p < 0.01, ***p < 0.001.

We measured the effects of shNPAS4 and cocaine conditioning on functional glutamatergic synapses on D1- and D2-MSNs by measuring spontaneous excitatory postsynaptic currents (sEPSCs), which assesses the combined excitatory synapses onto MSNs (see methods; Figure S7C-F). Cocaine conditioning and/or shNPAS4 in D1-MSNs produced no significant changes in sEPSC frequency or amplitude (Figure S7C,E). Similar to prior literature on the effects of abused substances on D2-MSNs ^20,56,70,71^, we observed that cocaine conditioning reduced sEPSC frequency (Figure S7D), but not amplitude (Figure S7F), in control D2-MSNs (black bars), and this reduction was abolished by shNPAS4 (Figure S7D). These data suggest that NPAS4 plays an important role in regulating drug-induced changes in excitatory synaptic transmission on D2-MSNs, but multiple glutamatergic inputs activate NAc MSNs and influence drug seeking ^13,72,73^.

One of the strongest glutamatergic inputs onto NAc D2-MSNs comes from the mPFC ^72,74^, and prelimbic (PrL) mPFC pyramidal neuron projections to the NAcore are required for cue-reinstated cocaine seeking in rats ^1,8,10,12,75-77^. To test the importance of the PrL→NAc circuit for mouse cocaine CPP, we infused in NAc a retrograde virus expressing Cre-recombinase (rgAAV-Cre) and a Cre-dependent Gi-DREADD in the mPFC (AAV2-hM4Di-mCherry) to allow for CNO-dependent suppression of PrL→NAc projection neuron activity during cocaine conditioning (Figure 6D; Figure S7B). We observed that Gi-DREADD-mediated suppression of the PrL→NAc circuit during cocaine conditioning significantly reduced cocaine CPP (Figure 6E, left). Gi-DREADD and CNO had no effects on total distance traveled (data not shown), and subsequent rounds of cocaine conditioning, in the absence of CNO, allowed for the development of normal place preference behavior (Figure 6E, right).

Since PrL→NAc circuit is important for both cocaine CPP and cue-reinstated cocaine seeking, we examined the PrL-specific synaptic transmission onto NAc D1- or D2-MSNs. To this end, we combined NAc-specific virus injections of Cre-dependent shNPAS4 in D1-Cre or D2-Cre mice with PrL-targeted virus injections of channelrhodopsin (AAV-hSyn-hChR2(H134R)-eYFP), then subjected mice to our standard 2-day cocaine or saline CPP procedure (Figure 6F,G). The next day we generated and analyzed coronal, acute slice recordings of blue light-evoked excitatory postsynaptic currents (oEPSCs) in NAc D1- or D2-MSNs (Figures 6H,I). shNPAS4 in D1-MSNs produced a small, but significant, increase in PrL→NAc oEPSC amplitude that was independent of cocaine conditioning (i.e., main effect of shNPAS4, but no interaction) (Figure 6H). In contrast, reduction of NPAS4 in D2-MSNs produced a significant interaction between cocaine conditioning and shNPAS4 virus (Figure 6I). Posthoc analyses revealed a robust increase in PrL→D2-MSN glutamatergic synaptic transmission in animals that received both shNPAS4 virus and cocaine conditioning (Figure 6I). These data reveal that NPAS4 in NAc D2-MSNs functions to limit strengthening of PrL→NAc D2-MSN inputs produced by cocaine conditioning, which serves to maintain the important D1-MSN:D2-MSN activation imbalance critical for drug seeking (Figure 6J).

## Discussion

A fundamental function of the brain is to form enduring associations between salient experiences and environmental cues, and neuronal activity-dependent mechanisms are important for this process process ^61,78^. Environmental cues and reward associations are crucial in SUD since powerful drug-cue associations often trigger drug seeking and relapse ^79^ The mechanisms by which reward-linked cues produce and maintain a preponderant activation of D1-MSNs to support drug-seeking behavior remain unclear. Here we showed that the neuronal activity-regulated transcription factor, Neuronal PAS Domain Protein 4 (NPAS4), plays an essential, cell type-specific role in NAc to control the D1- and D2-MSN activation balance, consistently reported as necessary for drug-seeking behavior ^13,14,16,19,80^. Using a new *NPAS4*-TRAp2a mouse combined with chemogenetics, we found that an NPAS4-expressing subpopulation of NAc neurons is required for expression of cocaine conditioned place preference. Single-cell transcriptomic analyses and *in situ* approaches from drug-conditioned mice revealed that NPAS4 is expressed predominantly in NAc MSNs, and using cell type-specific molecular genetic approaches, we found that NPAS4 in NAc D2-MSNs, but not D1-MSNs, is required for drug-context associations and cue-reinstated cocaine-, but not sucrose-, seeking behavior. We also show that NPAS4 in NAc D2-MSNs influences the activation balance of D1-MSNs and D2-MSNs, at least in part, by suppressing cocaine experience-dependent increases in synapse density and mPFC excitatory drive onto D2-MSNs. Analysis of differential gene expression in NAc D2-MSNs reveals that NPAS4 and cocaine conditioning regulates multiple genes whose functions are linked to synapses, neuronal projections, and the cocaine-regulated transcriptome, positioning NPAS4 as a critical regulator of D2-MSN synaptic connectivity and plasticity. Together our data reveal that NAc NPAS4-expressing neurons form a functional ensemble of neurons necessary for drug-cue associations, and that NPAS4 is a critical molecular player that allows addictive drugs to highjack brain reward circuits controlling prepotent, relapse-promoting drug-cue associations.

The activity balance between NAc D1-MSNs and D2-MSNs influences several drug-related behaviors, with D1-MSN activation typically promoting drug behavior and D2-MSNs opposing responses ^13,16,20,57^. Similar to drug-induced expression of FOS ^60,81^, we find that cocaine conditioning induces more NPAS4-positive NAc D1-MSNs than NAc D2-MSNs. The preponderant D1-MSN activation is likely produced by the well-studied influences of dopamine on D1-like receptor activation on glutamatergic synaptic transmission, the activity-dampening effects of D2-like receptor activation, and the known glutamatergic input differences onto these two NAc cell populations ^20,72,82^. We find that NPAS4 is induced in both NAc D1- and D2-MSNs during cocaine conditioning, but its cocaine-specific behavioral functions seem selectively required in D2-MSNs. Of note, viral-mediated reduction of NPAS4 in D1-MSNs significantly enhanced PrL→NAc D1-MSN glutamatergic synaptic transmission, but the effect was independent of cocaine conditioning, and perhaps basal activity states in D1-MSNs, but not D2-MSNs, are permissive for PrL input strengthening sans NPAS4. NPAS4 knockdown in NAc D1-MSNs did not alter cocaine CPP or cue-reinstated cocaine seeking, but since the cocaine doses were relatively high in these assays (i.e., 7.5 mg/kg and 0.5 mg/kg/infusion, respectively), we can’t rule out the possibility that NPAS4 knockdown in D1-MSNs might enhance cocaine-conditioned behaviors at low or subthreshold cocaine doses.

One of the principal targets of NAc MSNs is the dlVP, and D2-MSNs make strong connections onto these neurons ^26,62^. Optogenetic activation of NAc D2-MSNs that project to the dlVP attenuates contextual cocaine seeking ^9,26,83^, so NPAS4-dependent suppression of mPFC→NAc D2-MSN circuit strengthening is clearly important for place conditioning and drug-cue associations that promote cocaine seeking. Clearly, there are multiple glutamatergic inputs to the NAc that are important for regulating addiction-related behaviors in rodents ^13,73^, but the PrL→NAcore plays an essential role in cue-reinstated drug seeking ^10,12^, and we showed here that the activity of this same circuit is required for development of cocaine CPP. Compared to NAc D1-MSNs, NAc D2-MSNs receive a stronger excitatory input from the mPFC ^72^, suggesting the important need for NPAS4 to suppress drug-induced strengthening of glutamatergic inputs onto D2-MSNs. While we focused here on the behaviorally required mPFC→NAc circuit, it will be interesting in the future to explore the influence of NPAS4 on other major glutamatergic inputs onto NAc D2-MSNs, including basolateral amygdala and ventral hippocampus ^13^. Similar to a prior report ^56^, the sEPSCs in D2-MSNs, which measures combined excitatory inputs from all sources, showed a significant decrease in frequency, but not amplitude, following cocaine conditioning. Reduction of NPAS4 in D2-MSNs blocks the cocaine conditioning-induced decrease in sEPSC frequency, which might reflect the net balance of a reduction of non-mPFC inputs and the increase in mPFC→NAc D2-MSNs inputs, and/or an important role for NPAS4 in the reduction of non-mPFC synaptic inputs, which will be worth investigating in future studies.

There are likely multiple NPAS4 target genes (direct or indirect) in D2-MSNs which block dendritic spine formation and mPFC input strengthening produced by cocaine conditioning. We noted several interesting upregulated DEGs, including *Shisa9* and *Cartpt*. Shisa9, initially named CKAMP44, is involved in regulating short-term neuronal plasticity by interacting with AMPA receptors at dendritic spines to promote excitatory synaptic AMPA receptor desensitization ^65^. Cartpt (cocaine and amphetamine regulated transcript peptide) is a psychostimulant-regulated neuropeptide that appears to limit cocaine CPP ^66^. Another interesting DEG is *Penk*, a gene that encodes the reward-related neuropeptide, enkephalin, which binds to delta and mu opioid receptors with high affinity and is thought to limit GABA release by activation of DOR and MOR autoreceptor Gi-coupled signaling ^84,85^. In the NAc, DORs expressed postsynaptically on D2-MSNs ^86^ and on the terminals of PFC afferents ^87^. In control mice, cocaine conditioning significantly reduced *Penk* gene expression in D2-MSNs, and this was accompanied by a decrease in sEPSC frequency. However, the reduction in Penk mRNA was absent in cocaine-conditioned mice expressing NAc shNPAS4, revealing a novel role for NPAS4 in *Penk* expression in NAc D2-MSNs. Calmodulin genes are also differentially-expressed following cocaine conditioning (e.g., downregulated *Calm2* in S-shScram vs C-shScram) and shNPAS4 (upregulated Calm1 in C-shScram vs C-shNPAS4), which are involved in brain calcium signaling and differentially expressed in mice following cocaine SA ^88^. Interestingly, in humans who abused cocaine, there is a reduction in calmodulin-related gene transcription ^89^. It’s worth noting that NPAS4 mRNA and protein are very short-lived (∼60-90 minutes ^33,90^). As such, we speculate that some of the D2-MSN DEGs at 24-hrs post-cocaine conditioning are indirect targets of NPAS4. We chose to analyze this time point since it corresponds to the precise time when behavioral and synaptic changes are observed. A large fraction of the shNPAS4 D2-MSN DEGs are upregulated, and since NPAS4 typically increases expression of its target genes, many of these upregulated DEGs could be produced by secondary waves of gene expression (i.e., late response genes), produced as a consequence of synaptic changes influenced by shNPAS4 and cocaine conditioning, or reflect a potential repressor function for NPAS4 on some gene targets. Future studies examining multiple time points after cocaine conditioning will be important to understand the cascade of transcriptional events following cocaine conditioning. Finally, the largest number of cocaine-induced DEGs are found in the non-neuronal cells, including astrocytes and oligodendrocytes, and most of those DEGs are not influenced by NPAS4, which is strictly neuronal. Astrocytes play an important role in shaping synaptic transmission associated with drug-cue reactivity ^91^, and the cocaine conditioning-induced changes in glial gene expression highlight the strong engagement of these non-neuronal populations in drug-related behavior.

Taken together, our findings here reveal a novel role for NPAS4 in D2-MSNs to allow cocaine, but not sucrose, to form prepotent drug-cue associations and support relapse-like behavior. This is accomplished, at least in part, by blocking the cocaine experience-dependent strengthening of glutamatergic synaptic drive onto D2-MSNs, particularly from behaviorally required mPFC inputs, thereby maintaining the preponderant activation of D1-MSNs by drug-associated contexts and cues. NPAS4 is induced predominantly in NAc D1- and D2-MSNs following during place conditioning, and we show that the ensemble of NPAS4-inducing neurons during cocaine conditioning is essential for forming drug-context memories. NAc MSNs are functionally hyperpolarized and receive significant tonic inhibitory transmission from local interneurons under basal conditions ^7^, therefore, induction of NPAS4 in D2-MSNs plays an essential homeostatic role within these neurons to block synaptic plasticity that would otherwise shift the D1-MSN:D2-MSN activation balance and oppose the ability of drug-associated cues to trigger drug seeking. Understanding the mechanisms by which NPAS4 suppresses anti-relapse synaptic adaptations might reveal potential therapeutic avenues to block or reverse prepotent drug-cue associations that often trigger relapse in individuals recovering from SUD.

## Supporting information

Marker genes statistics for all clusters in the snRNA-seq datasets related to Figure 2.

Differential expression statistic for each comparison related to Figures 2 and 5.

## Acknowledgments

This work was supported by NIH grants F31 DA048557 (BWH), T32 DA07288 (BWH), NIH R01 DA032708 (CWC), and P50 Center on Opioid and Cocaine Addiction DA046373 (CWC). We would like to thank J. McGinty, P. Kalivas, J. Otis, E. Schmidt, T. Jhou, J. Woodward, C. Robinson, and S. Duncan for feedback on various aspects of the research designs and data analyses during my dissertation research. We would also like to thank M. Greenberg for sharing their anti-NPAS4 antibody. Some graphics and experimental design schema were created with Biorender.com. The opinions expressed in this article are the authors’ own and do not reflect the views of the NIH/NIDA.

## Materials and Methods

### Recombinant plasmids and shRNA expression viral vectors

For knockdown of endogenous NPAS4 mRNA expression in the NAc, previously validated NPAS4 shRNA (shNPAS4) or scramble (shScram) shRNA control was cloned into the pAAV-shRNA vector as previously described (Lin et al., 2008; Taniguchi et al., 2017). The adeno-associated virus serotype 2 (AAV2) vector consists of a CMV promoter driving eGFP with a SV40 polyadenylation signal, followed downstream by a U6 RNA polymerase III promoter and NPAS4 shRNA or scrambled (SC) shRNA oligonucleotides, then a polymerase III termination signal - all flanked by AAV2 inverted terminal repeats. For cell type-specific NPAS4 knockdown, NPAS4 shRNA or scrambled shRNA control was cloned into the pAAV-Sico vector (Ventura 2004 PNAS) with slight modifications. The Cre-dependent NPAS4 shRNA viral vector has a constitutively-active promoter driving expression of mCherry, and a second, LoxP site flanked eGFP expression cassette that physically separates the U6 promoter from the NPAS4 shRNA coding sequence. In the absence of Cre recombinase, the viral-transduced cells express both mCherry and eGFP (but no shRNA). However, when this virus enters a cell that expresses Cre recombinase (e.g. D1-Cre animal), the eGFP expression cassette is excised and the cell expresses only mCherry and the NPAS4 shRNA (no eGFP). AAV2-NPAS4 shRNA, Cre-dependent NPAS4 shRNA, and scrambled control shRNAs were packaged by USC Vector Core (Columbia, SC).

### Animals

All experiments were conducted in male and female animals 8-16 weeks of age during their dark phase. Targeted recombination in activated populations (TRAP) was achieved through design of a new “NPAS4-TRAP” mouse with NPAS4-dependent expression of cyclic recombinase that is fused with a triple mutant form of the human estrogen receptor (Cre-ERT2). Using these mice, NPAS4 is expressed along with Cre-ERT2, with a p2A cleavage site between that prevents changes in the function of endogenous NPAS4. When injected with a Cre-dependent AAV, such as DIO-mCherry, cells expressing NPAS4 will be labeled with mCherry following the injection of 4-hydroxytamoxifen (4OHT), which binds to ERT2 and allows nuclear translocation and recombination of floxed genes by Cre.

Other experimental mice included C57BL/6J (wild type, P60–80), heterozygous bacterial artificial chromosome (BAC) transgenic mice in which cyclic recombinase (Cre) was expressed under the control of the D1R promoter (Drd1a (D1)-Cre, line FK150) or the D2R promoter (Drd2 (D2)-Cre, line ER44), and BAC transgenic mice expressing tdTomato via the D1R promoter (D1-tdTomato) and eGFP via the D2R promoter (D2-eGFP). Mice were obtained from P. Kalivas (Medical University of South Carolina) and NINDS/GENSAT (http://www.gensat.org) (Gong 2007 J Neuro; Gong 2003 Nature). Rats expressing Cre under the control of the D1R (D1-Cre) or D2R promoter (D2-Cre) were obtained NIDA {Garcia-Keller, 2020 #520}, who assist us in validation of Cre expression in D1- and D2-expressing MSNs, respectively, then backcrossed to Charles River Long-Evans rats. Wild-type littermates were used for experimental controls. Both males and females were used for all experiments, and all animals were fed ad libitum, unless specified otherwise, and maintained at ∼22 °C in a humidity-controlled environment on a reverse 12-h light/dark cycle (lights off at 8:00 a.m.). Animals were single-housed after surgery. All experiments were conducted in accordance with the US National Institutes of Health Guidelines for the Care and Use of Laboratory Animals and all procedures were approved by the Institutional Animal Care and Use Committee at the Medical University of South Carolina.

### Viral-Mediated Gene Transfer

Stereotaxic surgery was performed under general anesthesia with isoflurane (induction 4-5% v/v, maintenance 1%-2% v/v). Coordinates to target the NAc (mainly core subregion) were +1.6 mm anterior, +1.5 mm lateral, and −4.4 mm ventral from bregma (relative to skull) at a 10 degree angle in mice, and +1.6 mm anterior, +2.8mm lateral and −7.3 mm ventral (relative to skull) in rat. Coordinates to target the mouse mPFC prelimbic cortex were +1.85 mm anterior, +0.75 mm lateral, and −2.3mm ventral from bregma at a 15 degree angle. AAV-NPAS4 shRNA, AAV-Cre-dependent NPAS4 shRNA, AAV-DIO-mCherry, AAV-DIO-GiDREADD-mCherry, AAV-Retro-Cre, and scrambled control shRNAs were delivered using Hamilton syringes or Nanoject III at a rate of 0.1 uL/min for a total 0.4-0.6 uL/hemisphere in mice and a total 0.6-0.8 uL/hemisphere in rats, with an additional 7-10 min before needles were retracted, as previously described (Taniguchi et al., 2012, Taniguchi et al., 2017). Viral placements were confirmed by AAV-mediated fluorophore expression (i.e. GFP or mCherry) using confocal microscopy.

### Immunohistochemistry

Mouse brains were drop-fixed overnight in 4% PFA in 1X PBS and transferred to a 30% sucrose solution in 1X PBS before slicing (40 or 50 mm) on a sliding microtome (Leica Microsystems). The slices were permeabilized and blocked in 3% BSA, 0.3% Triton X-100, 0.2% Tween-20, 3% normal goat serum in PBS, then incubated with primary antibodies: anti-NPAS4 (1:1000, kindly provided by Dr. Michael Greenberg’s lab), anti-mCherry (1:500, chicken, LSBio), or anti-FOS (1:1000, rabbit, Synaptic Systems) in blocking buffer at room temperature for 2 hr. Following a series of 1X PBS rinses, slices were incubated for 2 hr at room temperature with secondary antibodies (donkey anti-rabbit 488, donkey anti-rabbit 594, or donkey anti-Chicken 594) while protected from light. Slices were counterstaining with Hoechst, mounted with ProLong Gold, coverslipped on glass slides, then analyzed with confocal microscopy (Zeiss LSM 880 or Leica Stellaris). Protein expression measured using ImageJ software under experimenter blinded conditions.

### Fluorescent in situ hybridization

Mice were live decapitated, then the brains were rapidly extracted and flash frozen in isopentane solution on dry ice. Brains were sliced at 16 µm in a cryostat (−18°C), and sections were air-dried to the slides at −18°C and stored at −80°C. For the detection of D1 receptor (Drd1a), D2 receptor (Drd2), Cre recombinase, or NPAS4 mRNA, the RNAscope Fluorescent Multiplex kit (320293; Advanced Cell Diagnostics) was used with slight modification of manufacturer instructions and using commercially available probes [NPAS4 (C1, #423431), iCre (C2, #423321), Drd1a (C3, #406491), and Drd2 (C3, #406501)], as described previously (Garcia-Keller, 2020, J Neuro). Briefly, sections were fixed for 20 min in neutral buffered 10% formalin, followed by ethanol dehydration series on 50%, 70%, 95%, and 100%, 5 min each. Before hybridization sections were treated with protease IV (322340, Advanced Cell Diagnostics) for 10 minutes. Sections were hybridized for 2 h at 40°C, and probes were amplified using TSA fluorescent detection kits as directed by manufacture (Akoya Bioscience). The C1 probe was conjugated to Fluorescein, C2 to Cy3, and C3 to Cy5. Sections were then counterstained with DAPI and coverslipped using ProLong Gold mounting medium (Thermo Fisher) and allowed to dry. Slices were then imaged via confocal microscopy (Zeiss LSM 880).

### Cocaine conditioned place preference

Mice were conditioned to cocaine using an unbiased paradigm described previously (Smith et al., 2016; Taniguchi et al., 2017). On day 1, mice were placed in the chamber and allowed to explore the conditioning apparatus, which consisted of 2 distinct environment chambers (white side or black side with different texture flooring). Mice that showed pre-conditioning preference (more than 30% of total time spent in either of the 2 chambers) were excluded from the study. On days 2 and 3, mice received cocaine (7.5 mg/kg; i.p.) in the AM and were confined to one chamber. In the PM sessions on days 2 and 3, 5-6 hours after AM conditioning, mice received saline (0.9%; 1 ml/kg; i.p.) and were confined to the opposite chamber. On day 4, mice were placed again in one side of the CPP apparatus with free access to both chambers, and the time spent in each side was quantified. C57Bl/6J mice that were infused with AAV-Retro Cre in the NAc core and AAV-Cre-dependent Gi-DREADD mCherry in the PrL were subjected to a 6-day CPP paradigm, with cocaine pairings (7.5 mg/kg; i.p.) on days 2 and 4, and saline on days 3 and 5 on the opposite side of the place preference chamber. On the post-test (day 6), the mice were placed again in the CPP chamber with free access to both sides, and the time spent in each side was quantified. Data are expressed as time spent on the cocaine-paired side minus the time spent on the saline-paired side (CPP score) during the post-test.

### Locomotor Sensitization

Locomotor activity was measured via photobeam array (San Diego Instruments) before and after each injection using the two-injection protocol of sensitization (Valiente and Marín, 2010; Taniguchi et al., 2017). Mice received saline injections for 3 days, then on the 4th day, mice received an injection of cocaine (20 mg/kg; i.p.). After 5 days, the mice received a 2nd cocaine injection (20 mg/kg; i.p.) in the locomotor apparatus. Data are shown as the sum of total beam breaks in the first 30 min of each session.

### Rodent Cocaine or Sucrose Self-administration

Following Cre-dependent NPAS4 shRNA or scrambled shRNA infusion into the NAc (described above), rats were allowed to recover for 1 week prior to catheter implantation. Rat SA sessions (2 hr) were performed during the light phase at the same time each day, during which rats were placed in an operant conditioning chamber and connected to a drug line controlled by an external delivery pump. A ‘‘house’’ light inside the chamber signaled drug availability. All chambers contained an active (50 uL, 3 s infusion; 0.5 mg/kg cocaine) and an inactive lever. During an infusion, a cue light above the lever was illuminated and followed by a 20 s time-out period signaled by the house light going off. Rats self-administered cocaine on a fixed ratio (FR) schedule beginning at FR1, followed by an FR3 and finally an FR5. When increasing the FR requirements, intake was analyzed for stability (< 20% variability within the last 3 sessions). Rats that completed 15-20 days of FR training and were stable on the FR5 schedule for at least 3 days continued to subsequent studies. Self-administration sessions (2hr) were performed at the same time each day during the dark phase as described previously (Bobadilla et al., 2017). Briefly, drug or sucrose availability was signaled both by the house light and a light above the active nose poke hole. Following a poke in the active hole both availability lights went off and a cue light inside the nose poke hole was illuminated. Cocaine (12 ul, 2 s infusion; 0.5 mg/kg,) or a sucrose pellet (15 mg) were delivered immediately upon the active nosepoke, followed by a 10 s time-out period. Nose pokes in the inactive hole were without programmed consequences. Rats that met requirements for stable cocaine self-administration (last 3 days > 85% discrimination between active and inactive nose poke hole and > 10 infusions) entered into a 7-day abstinence phase.

### Extinction and Reinstatement

Rats were allowed a week of forced abstinence (withdrawal) undisturbed in their home cages. Then, animals were placed back into the operant chambers and lever pressing in the absence of any drug administration, cues, and timeouts was measured on both levers. Extinction training continued for at least 6 days, followed by reinstatement sessions. Each reinstatement session consisted of a 2 hr extinction session. Priming stimuli included presentation of the drug paired cue (light) or experimenter-administered cocaine (10 mg/kg, i.p.). Lever pressing was measured during the 2 hr session. During cue-induced reinstatement, reward availability (house and active port light) was returned as active lever pressing output, but drug or sucrose delivery was omitted.

### Dendritic spine morphometric analyses

D2-Cre mice received bilateral NAc injections of our Cre-dependent shNPAS4 mixed 1:1 with a Cre-dependent spaghetti monster FLAG virus (AAV-CAG-Ruby2sm-FLAG-WPRE-SV40) to label D2-MSN dendritic spines using anti-FLAG IHC. Brains were collected after perfusion with 4% PFA in 1x PB, 24 hr after the last cocaine conditioning session, and fixed overnight in 4% PFA in 1x PB, then transferred to a 30% sucrose solution in 1x PB before slicing (100 um) with a vibratome. D2-MSNs (expressing mCherry and 647-labeled FLAG) were sampled for dendritic spine analyses as described previously (Siemsen et al., 2019). Briefly, dendrites past the secondary branch point and ≥ 100 um from the cell body were imaged with a Leica SP8 laser scanning confocal microscope equipped with HyD detectors for enhanced sensitivity. Dendritic spine segments were selected only if they satisfied the following criteria: 1) could clearly be traced back to a cell body of origin, 2) were not obfuscated by other dendrites, and 3) were co-labeled with somatic mCherry, but not eGFP, ensuring spines were sampled only in recombined, Cre+ neurons. Images were collected with a 63X oil immersion objective (1.4 N.A.) at 1024×512 frame size, 4.1X digital zoom, and a 0.1µm Z-step size (0.04×0.04×0.1 µm voxel size). Pinhole was set at 0.8 airy units and held constant. Laser power and gain were empirically determined and then held relatively constant, only adjusting to avoid saturated voxels. Huygens Software (Scientific Volume Imaging, Hilversum NL) was used to deconvolve 3D Z-stacks. Deconvolved Z-stacks were then imported into Imaris (version 9.0.1) software (Bitplane, Zurich CH). The filament tool was then used to trace and assign the dendrite shaft. Dendritic spines were then semi-automatically traced using the autopath function, and an automatic threshold was used to determine dendritic spine head diameter. Variables exported included the average spine head diameter (in µm) as well as the number of dendritic spines per µm of dendrite (spine density). 3-10 segments were sampled per animal, and the average spine head diameter and the spine density were calculated for each segment. Data for each variable was then expressed as number of spine segments/number of animals. All analyses were performed under experimenter-blinded conditions.

### Electrophysiology

All acute-slice electrophysiological experiments were performed in shScram and shNPAS4 D1- or D2-Cre mice at 11-13 weeks old. Acute coronal slices (300-μm thickness) containing mPFC and NAc were prepared in a semi-frozen 300 mOsM dissection solution containing (in mM): 100.0 choline chloride, 2.5 KCl, 1.25 Na2H2PO4, 25.0 NaHCO3, 25.0 D-glucose, 3.1 Na-pyruvate, 9.0 Na-ascorbate, 7.0 MgCl2, 0.5 CaCl2 and 5.0 kynurenic acid (pH 7.4) and was continually equilibrated with 95% O2 and 5% CO2 prior to and during the slicing procedure. Slices were transferred to a 315 mOsM normal artificial cerebrospinal fluid (ACSF) solution containing (in mM): 127 NaCl, 2.5 KCl, 1.20 Na2H2PO4, 24 NaHCO3, 11 D-glucose, 1.2 MgCl2, 2.40 CaCl2, and 0.4 Na-ascorbate (pH 7.4) to recover at 37°C for 30 minutes, and then transferred to room temperature ACSF for an additional 30 minutes prior to recording. NAc MSNs (depth 30-100 μm into the slice) were visualized with infrared differential interference contrast optics (DIC/infrared optics) and identified by their location, apical dendrites, and burst spiking patterns in response to depolarizing current injection. Cre-expressing cells (D1- or D2-MSNs) were identified by Cre-dependent viral expression of mCherry and the absence of GFP (which is removed in Cre-expressing cells to induce shRNA expression). Unless stated otherwise, all electrophysiological experiments were performed in whole cell voltage clamp mode at −70 mV using borosilicate pipettes (4-6 MΩ) made on NARISHIGE puller (NARISHIGE, PG10) from borosilicate tubing (Sutter Instruments) and filled by an internal solution containing (in mM): 140.0 CsMetSO4, 5.0 KCl, 1 MgCl2, 0.1 CaCl2, 0.2 EGTA, 11 HEPES, 2 NaATP, 0.2 Na2GTP (pH 7.2; 290– 295 mOsm). All data (Recordings) were acquired and analyzed by amplifier AXOPATCH 200B (Axon Instruments), digitizer BNC2090 (National instruments) and software AxoGraph v1.7.0, Clampfit v8.0 (pClamp, Molecular Devices) and MiniAnalysis Program v6.0.9 (Synaptosoft). Data were filtered at 2 kHz via AXOPATCH 200B amplifier (Axon Instruments) and digitized at 20 kHz via AxoGraph v1.7.0.

### Optically evoked postsynaptic currents

The evoked postsynaptic responses of NAc D1- or D2-MSNs were elicited by blue light stimulation of excitatory afferents from the mPFC as previously described (Deroche et al., 2020). A 473 nm laser (Dragon Laser) coupled to a 50 μm core glass silica optical fiber (ThorLabs) was positioned directly above the slice orientated 30° ∼350 μm from the recording electrode. At the site of recording discounting scattering a region of ∼0.05 mm2 was illuminated that after power attenuation due to adsorption and scattering in the tissue was calculated as ∼100 mW/mm2 (Yizhar et al., 2011; Deroche et al., 2020). Optically evoked EPSCs were obtained every 10 s with pulses of 473 nm wavelength light (0–10 mW, 2 ms). AMPA-receptor-mediated excitatory postsynaptic currents (EPSCs) were recorded in presence of picrotoxin (100 μM, Sigma Aldrich) to block GABAARs. Outliers were assessed by a Grubbs test (alpha=0.05) in GraphPad Prism software and excluded from analysis.

### Spontaneous postsynaptic currents

Excitatory spontaneous postsynaptic currents (sEPSCs) were recorded from NAc MSNs in voltage clamp mode at −70 mV. MSNs were identified by their morphology parameters (medium-sized and round-shape soma, spiny dendrites) and by bursting pattern of action potentials firing in response to depolarizing current injection. For recordings we chose specific Cre-expressing MSNs (D1- or D2) which were identified by Cre-dependent viral expression of mCherry and the absence of GFP (which is removed in Cre-expressing cells to induce shRNA expression). Data were recorded in a series of 10 traces (sweeps), 10 second each. At the beginning of each sweep, a depolarizing step (4 mV for 100 ms) was generated to monitor series (10-40 MΩ) and input resistance (>400 MΩ). To analyze data synaptic events were detected via custom parameters in MiniAnalysis software (Synaptosoft, Decatur, GA) and subsequently confirmed by observer. Data were measured until 700 events in a series were analyzed, or until the maximal duration of the series.

### Single nuclei RNA-seq and bioinformatic analysis

Wild-type C57BL/6J mice received bilateral NAc infusion of AAV-NPAS4 shRNA-GFP or scrambled-GFP control and underwent conditioned place preference for cocaine or saline (total of 4 groups with 4 animals/group). Mice were then rapidly decapitated at 12 weeks of age, and the brains extracted into 4°C Hibernate A medium with the GlutaMAX, B27 supplement, and NxGen RNase inhibitor (0.2U/uL; Lucigen). NAc tissues were dissected rapidly, flash frozen, and stored at −80°C until the day of dissociation. These specific steps were modified from Savell et al., 2020 by Hughes, B.W. The following day (when all samples were prepped), frozen tissues were slowly thawed on ice, then chopped with a razor blade ∼75-100 times in 2 orthogonal directions. Chopped NAc tissue was lysed with hypotonic lysis buffer (10mM Tris-HCL, 10mM NaCl, 3mM MgCl2, 0.1% (v/v) IGEPAL), then supplemented Hibernate-A medium was added, tissue triturated 15-20 times/sample using 3 glass capillaries with decreasing diameters, then 40-micron filtered. Nuclei were isolated by 500xg centrifugation, washed with 1X PBS + 1% BSA and 0.2U/uL RNase inhibitor. Nuclei were then incubated in 7AAD and underwent FACS to remove clumps and whole cells. Samples were then counted and diluted to 1500 nuclei/ul before immediate processing using the 10x Genomics Single-Cell Protocol by the MUSC Translation Science Lab. Libraries were constructed using the Chromium Single Cell 3’ Library Construction Kit (10x Genomics, v3.1), sequenced at Vanderbilt’s Next Gen Sequencing Core (Illumina NovaSeq 6000).

### Sequencing analysis

Raw sequencing data were processed with Cell Ranger (v6.1.2) (PMID: 28091601). Cellranger mkfastq command was used to demultiplex the different samples and cellranger count command was used to generate gene – cell expression matrices. Ambient RNA contamination was inferred and removed using CellBender (v0.232) with standard parameters. Mouse genome mm10 was used for the alignment and genecode vM25 was used for gene annotation and coordinates (PMID: 33270111).

Downstream analysis was performed in R with Seurat (v4.1.0) (PMID: 34062119) and customized R scripts. Data from different samples (S-scScram, C-scScram, S-scNPAS4, C-scNPAS4) were merged into a unique single cell object. Nuclei with > 250 genes, < 10000 UMIs, and < 5% mitochondrial transcripts were retained for downstream analysis. Genes located in the mitochondrial genome were removed. Doublets were removed using scDblFinder (v1.8.0) (PMID: 35814628) for a resultant expression matrix with 21,516 genes and 52,743 cells. The SCTransform workflow was used for count normalization (VST transformation) and data integration (PMID: 31870423) using 30 principal components and resolution of 0.5 for Louvain clustering and UMAP. Cluster marker genes were identified using FindAllMarkers using Wilcoxon Rank Sum test with the standard parameters. After removing 4 clusters showing high expression of glutamatergic neuronal markers and subsequent re-clustering, we identified 20 final distinct clusters. Cell annotation was performed using two different approaches: 1) markers enrichment using a Nucleus Accumbens independent data (PMID: 30257220) by Fisher’s exact test and 2) a predictive machine learning model based on a Nucleus Accumbens atlas reference data (PMID: 34663959) using the R package scPred (v1.9.2) (PMID: 31829268). Cell annotation then was manually curated to reflect both analyses.

### Differential expression analysis

To identify differentially expressed genes between conditions (S-shScram vs C-shScram, C-shScram vs C-shNPAS4) in each defined cell types, the R package LIBRA (v1.0.0) with option MAST was used to perform zero-inflated regression analysis (PMID: 34584091). Genes were defined as significantly differentially expressed at Benjamini–Hochberg correction FDR < 0.05 and abs(log2(Fold Change)) > 0.2. We further confirmed these results using a Wilcoxon’s rank sum test.

### Gene ontology analyses

The functional annotation of the identified DEGs was performed using ToppGene (PMID: 19465376). A Benjamini-Hochberg FDR (FDR<0.05) was applied as a multiple comparison adjustment. Enrichment was further confirmed using the R package clusterProfiler (v4.2.2) (PMID: 22455463).

### Data availability

All processed data, UMAP coordinates, and annotations have been made freely available to download and inspect at the BioCM portal (https://github.com/BioinformaticsMUSC/HughesEtAl_NPAS4Cocaine). User-friendly interactive data is available through ShinyCell at https://bioinformatics-musc.shinyapps.io/Hughes_NAc_Cocaine_NPAS4/. Raw and processed data to support the findings of this study have been deposited in GEO under accession number: GSE210850.

### Code availability

All code used to analyze the data and to generate the main figures of this paper can be found at https://github.com/BioinformaticsMUSC/HughesEtAl_NPAS4Cocaine/

### Statistics

Student t-tests and two-way or three-way analyses of variance (ANOVAs) with or without repeated-measures (RM) were used, with ANOVAs followed by Sidak or Tukey post hoc tests when a significant interaction was revealed, to analyze mRNA expression, NPAS4 signal intensity, cocaine conditioned place preference, cocaine and sucrose self-administration, electrophysiology, and dendritic spine morphometric data. All statistics were performed using GraphPad Prism, except single nuclei RNA-sequencing analysis that was performed in R. Statistical outliers were detected using a Grubbs test and excluded from analysis. All data are presented as the mean ± SEM. Significance was shown as # = p < 0.1, * = p < 0.05, ** = p < 0.01, *** = p < 0.001, **** = p < 0.0001, and non-significant values were either not noted or shown as n.s.

## Supplementary Information for

### Supplementary materials for this manuscript include the following

- **Figure S1**. Validation of NPAS4-TRAP mice, quantification of 4OHT-dependent labeling of NPAS4-expressing cells, and cocaine conditioning-dependent reactivation of endogenous NPAS4 after cocaine or saline conditioning TRAP.
- **Figure S2**. Single nuclei RNA-seq quality check and cluster comparison to previously published NAc data, as well as D2-MSN-specific DEGs between different group comparisons.
- **Figure S3**. Validation of Cre-dependent NPAS4 shRNA relative to scrambled shRNA control and locomotor sensitization following cell type-specific NPAS4 knockdown.
- **Figure S4**. No effects on acquisition, extinction, and cocaine-primed reinstatement following NPAS4 knockdown in D1- or D2-MSNs.
- **Figure S5**. NPAS4 knockdown in NAc D2-MSNs increases cue-induced reinstatement to natural reward seeking.
- **Figure S6**. Single nuclei RNA-sequencing of the mouse NAc after NPAS4 knockdown and cocaine CPP in D1-MSNs and comparison between D1- and D2-based clusters.
- **Figure S7**. Spine head diameter analysis in D2-MSNs, representative image of PrL→NAcore virus placements, and NPAS4 knockdown in D2-MSNs, but not D1-MSNs, affects spontaneous EPSC frequency.

### Supplementary tables (external attachments)

- **Supplementary Table 1:** Marker genes statistics for all clusters in the snRNA-seq datasets related to Figure 2.
- **Supplementary Table 2:** Differential expression statistic for each comparison related to Figures 2 and 5.

## Extended Data Figures S1-S7

**Extended Data Fig. 1:**
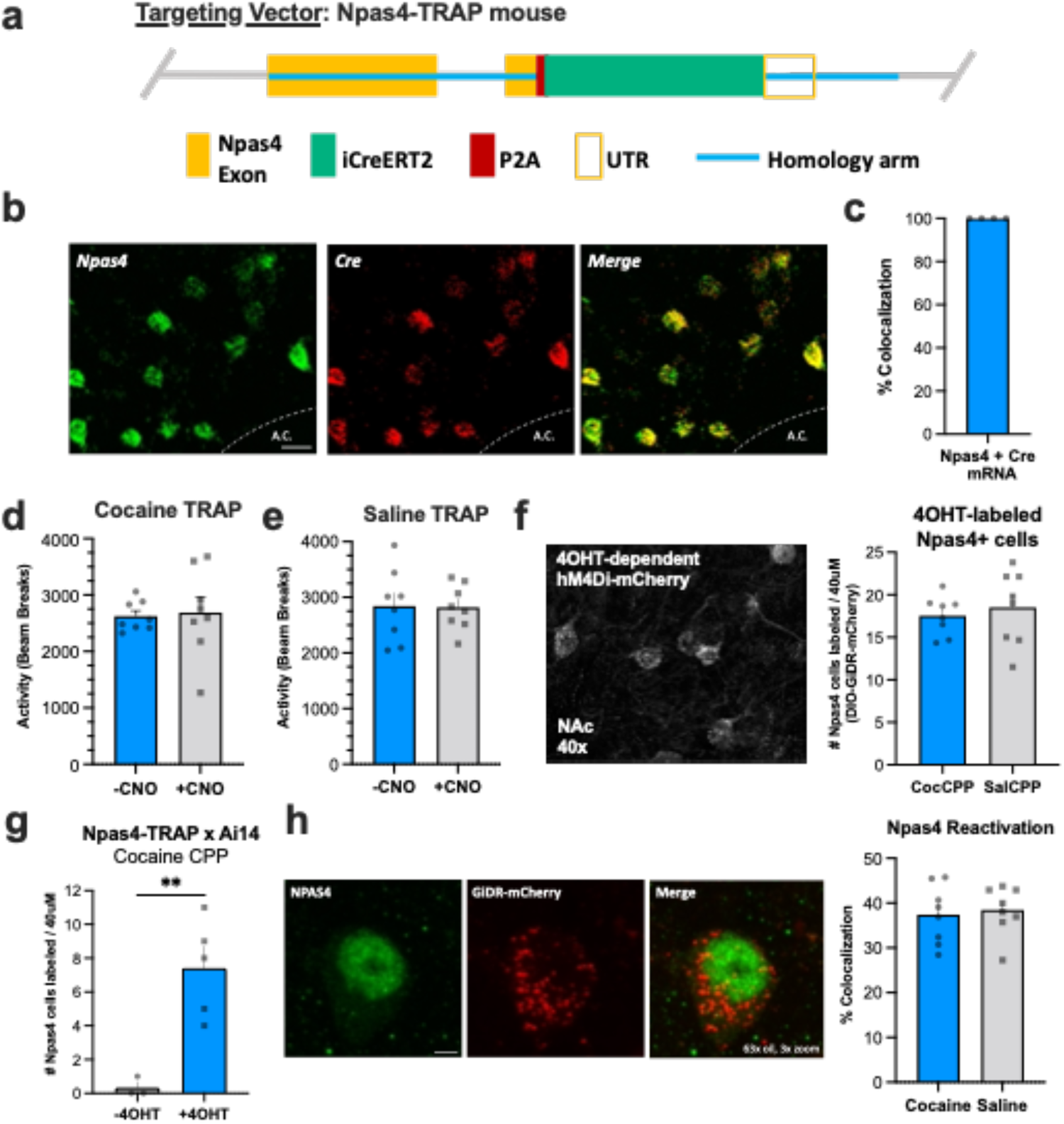
Validation of NPAS4-TRAP mice, quantification of 4OHT-dependent labeling of NPAS4-expressing cells, and cocaine/environment-dependent reactivation of endogenous NPAS4 after TRAP labeling during cocaine or saline conditioning. **a**, Targeting vector design for CRISPR/Cas9 generation of new NPAS4-TRAP mice. **b-c**, Representative image showing NPAS4 mRNA colocalization with Cre following cocaine CPP and validation of NPAS4-TRAP mouse. **d-e**, Total activity after 4OHT labeling in cocaine CPP and saline CPP, both showing no effect of post-test CNO on total locomotion. **f**, Representative image showing mCherry-labeled neurons that expressed NPAS4 during cocaine conditioning (left), and quantification of the number of cocaine- and saline-labeled cells by 4OHT (right). **g**, Number of NPAS4 TRAPed cells (Ai14) with or without 4OHT immediately after cocaine conditioning. **h**, IHC of cocaine or saline TRAPed neurons colocalized with cocaine CPP-induced NPAS4 reactivation and quantification after cocaine re-exposure (left) and reactivation of NPAS4-positive cells by a third cocaine conditioning session 4OHT after cocaine TRAP or saline TRAP (right). Data shown are mean ± SEM.

**Extended Data Fig. 2:**
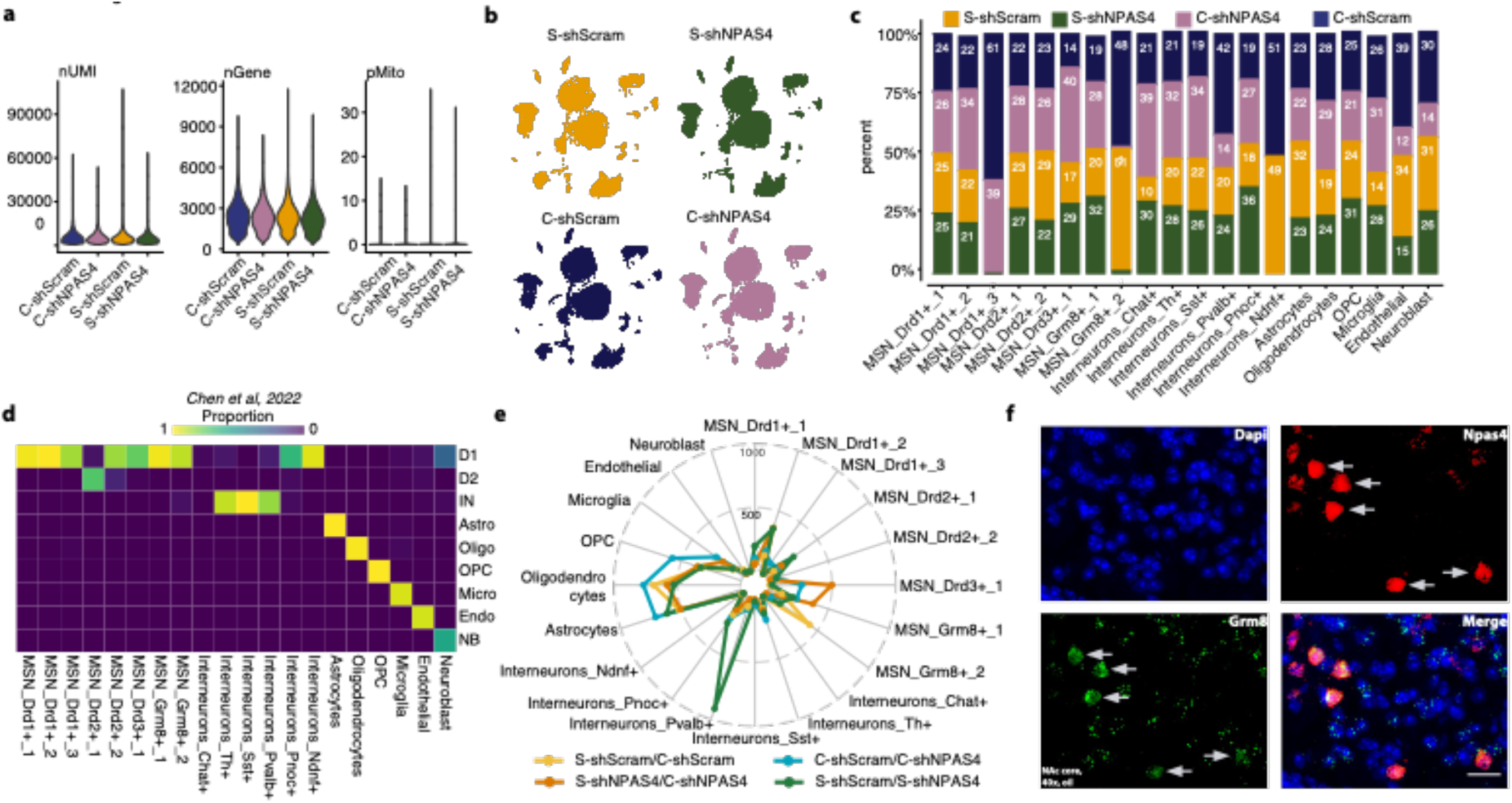
Single nuclei RNA-seq quality check and cluster comparison to previously published NAc data, D2-MSN-specific DEGs between different group comparisons, and confirmation of NPAS4 mRNA in Grm8-expressing MSNs. **a**, Violin plots showing the number of UMIs, total genes, and mitochondrial genes in each group. **b-c**, UMAPs showing similar cluster distribution between groups and similar numbers of each cell type in each group. **d**, Overlap of defined clusters from this paper compared to Chen et al., 2021. **e**, Radar plot showing the number of differentially expressed genes in each cluster with each group comparison. **f**, Fluorescent *in situ* hybridization confirming NPAS4 mRNA expression in NAc cells expressing Grm8 mRNA (i.e., NPAS4+ Grm8-MSNs).

**Extended Data Fig. 3:**
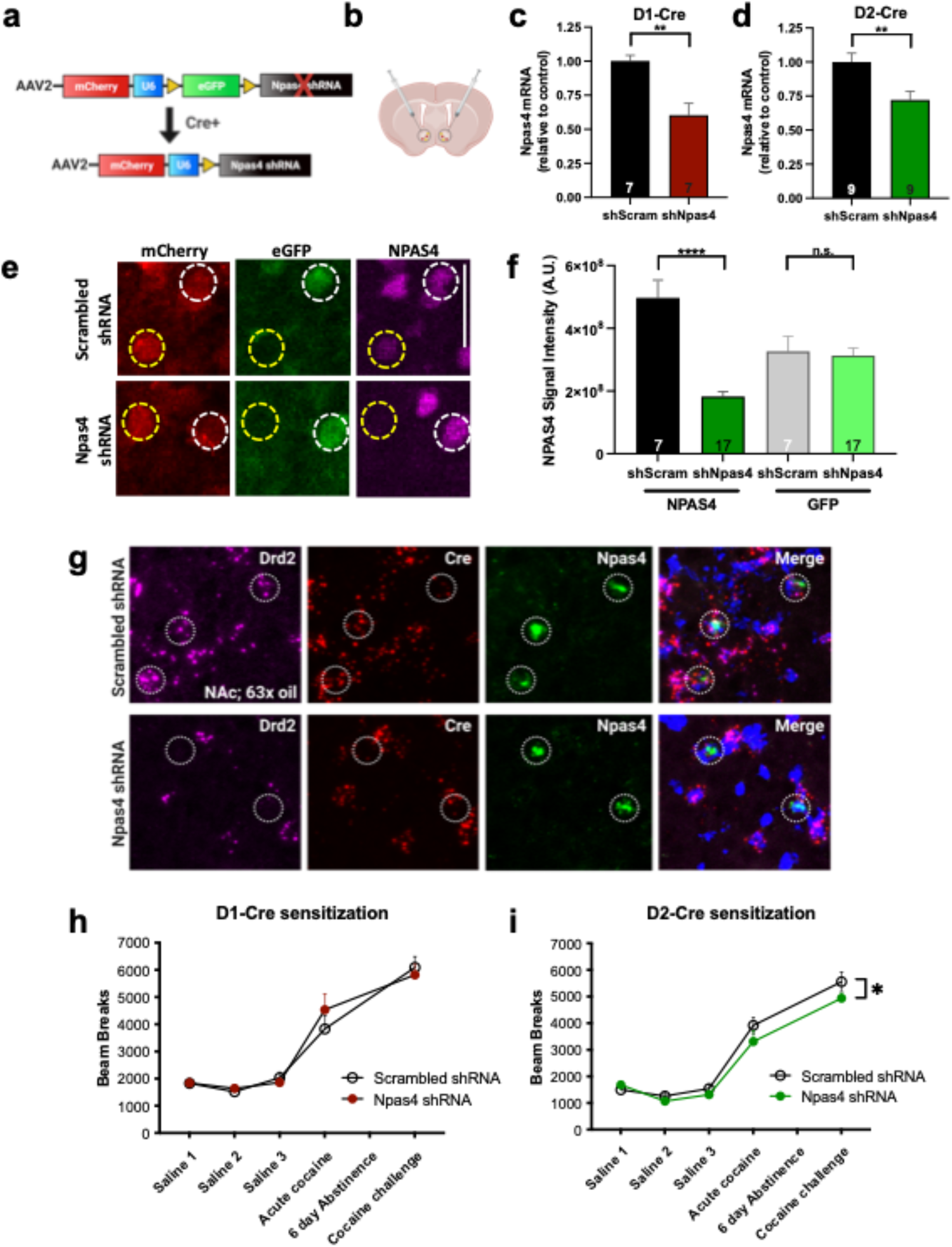
Validation of Cre-dependent NPAS4 shRNA relative to scrambled shRNA control and locomotor sensitization following cell type-specific NPAS4 knockdown. **a-b**, Vector design of Cre-dependent NPAS4 shRNA when infused into the NAc of a D2-Cre mouse. **c-d**, qPCR validation of NPAS4 knockdown in D1-MSNs of D1-Cre mice and D2-MSNs of D2-Cre mice. **e-f**, IHC showing Cre-dependent NPAS4 knockdown and quantification of NPAS4 knockdown and (right) no change in eGFP expression in shScram vs. shNPAS4. **g**, RNAscope showing Cre-dependent NPAS4 knockdown in D2-Cre mice. **h-i**, Locomotor sensitization following NPAS4 knockdown in D1-MSNs and D2-MSNs. Data shown are mean ± SEM; *p < 0.05, **p < 0.01, ****p < 0.0001, ns = not significant.

**Extended Data Fig. 4:**
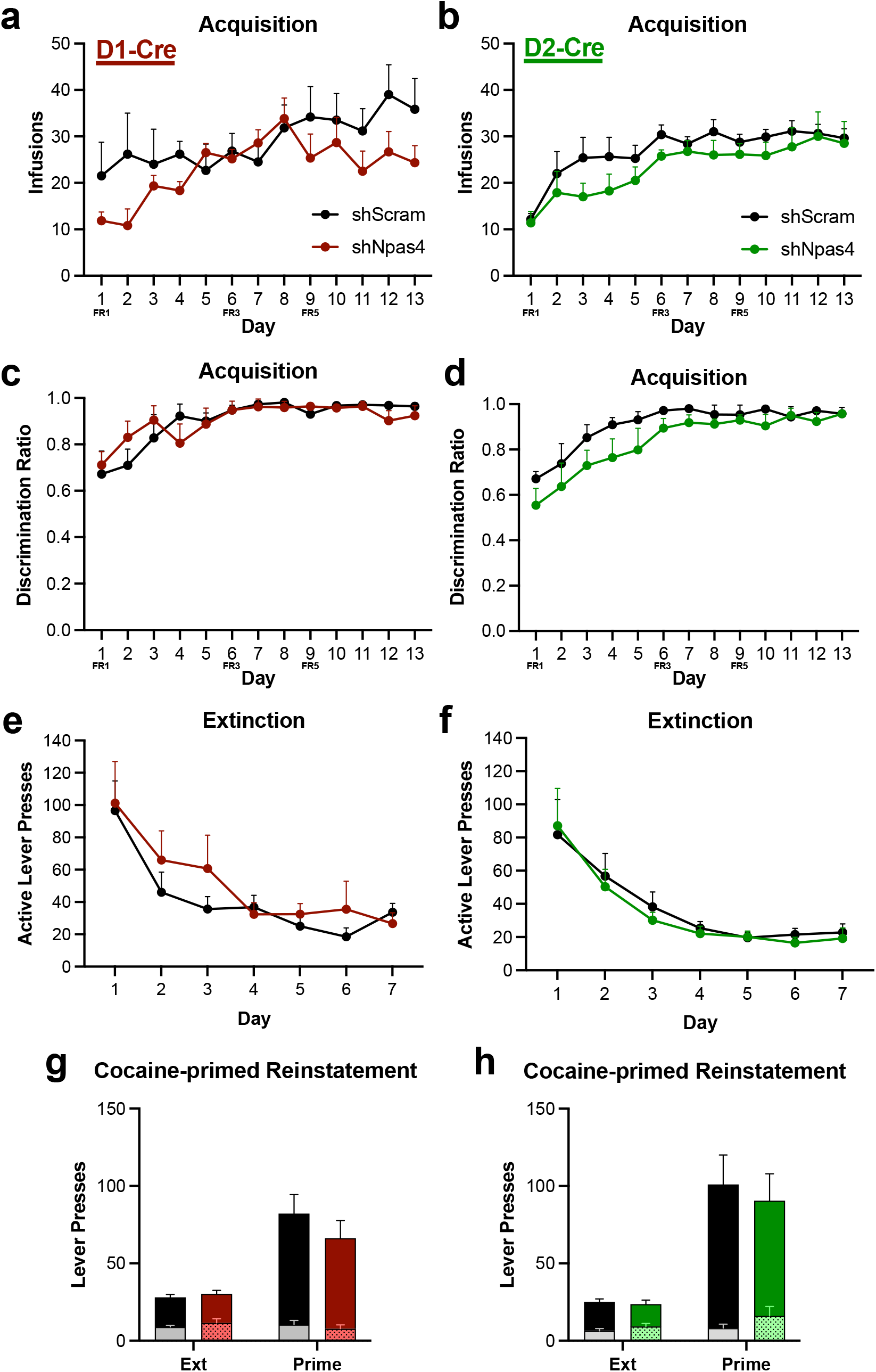
No effects on acquisition, extinction, and cocaine-primed reinstatement following NPAS4 knockdown in D1- or D2-MSNs. **a-b**, The number of infusions D1-Cre and D2-Cre rats received during cocaine SA acquisition following cell type-specific NPAS4 knockdown. **c-d**, Discrimination ratio between active and inactive presses during acquisition with NPAS4 knockdown in D1- and D2-MSNs. **e-f**, Active lever presses during extinction after NPAS4 knockdown in D1- and D2-MSNs. **g-h**, Cocaine-primed reinstatement after NPAS4 knockdown in D1- and D2-MSNs. Data shown are mean ± SEM.

**Extended Data Fig. 5:**
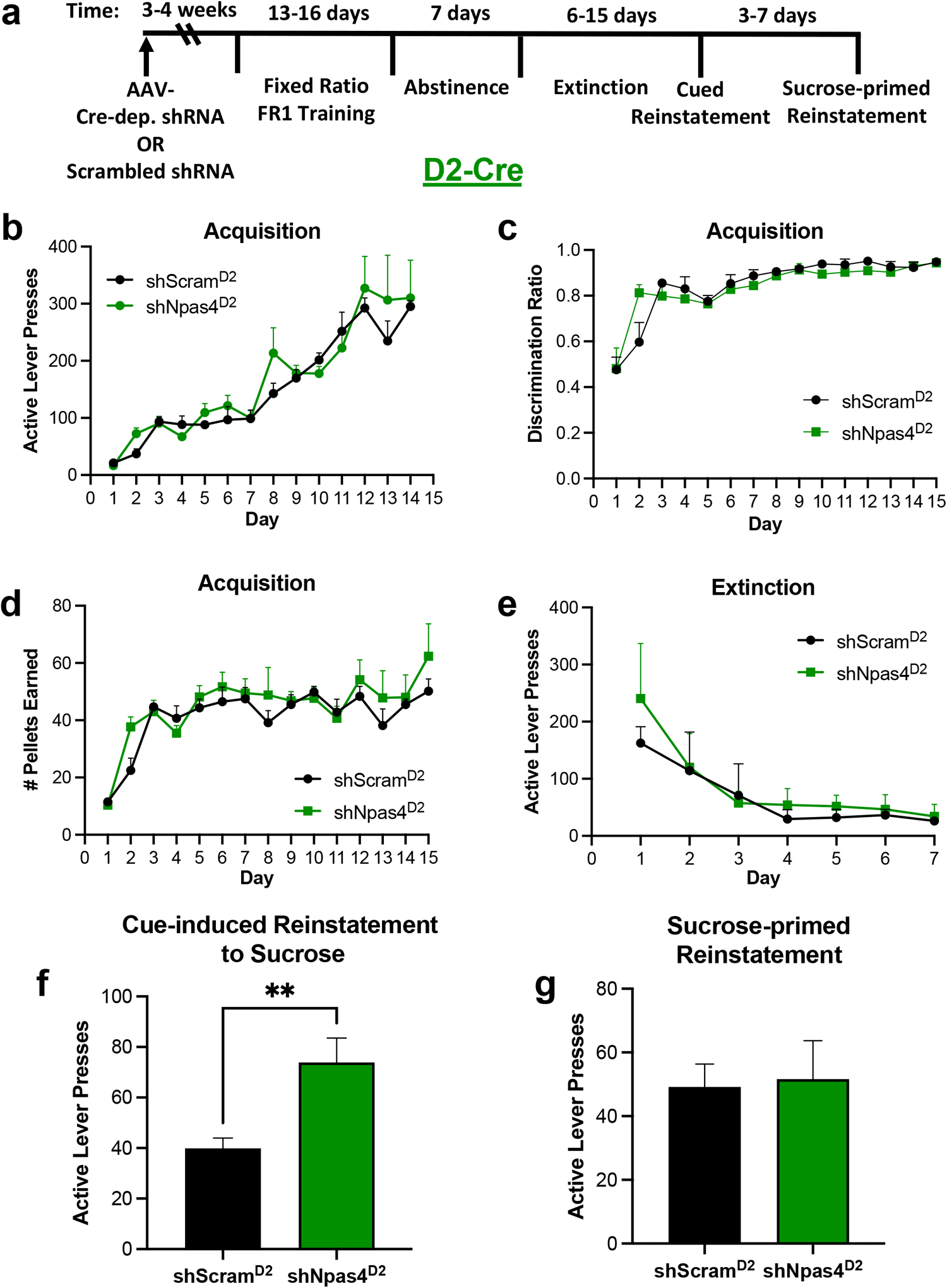
NPAS4 knockdown in NAc D2-MSNs increases cue-induced reinstatement to natural reward seeking. **a**, Timeline of sucrose SA after NPAS4 knockdown in D2-MSNs of D2-Cre rats. **b-e**, Active lever presses, discrimination ratio, and the number of pellets earned during acquisition, followed by extinction training. **f-g**, Cue-induced reinstatement to sucrose seeking and sucrose-primed reinstatement following NPAS4 knockdown in NAc D2-MSNs. Data shown are mean ± SEM; **p < 0.01.

**Extended Data Fig. 6:**
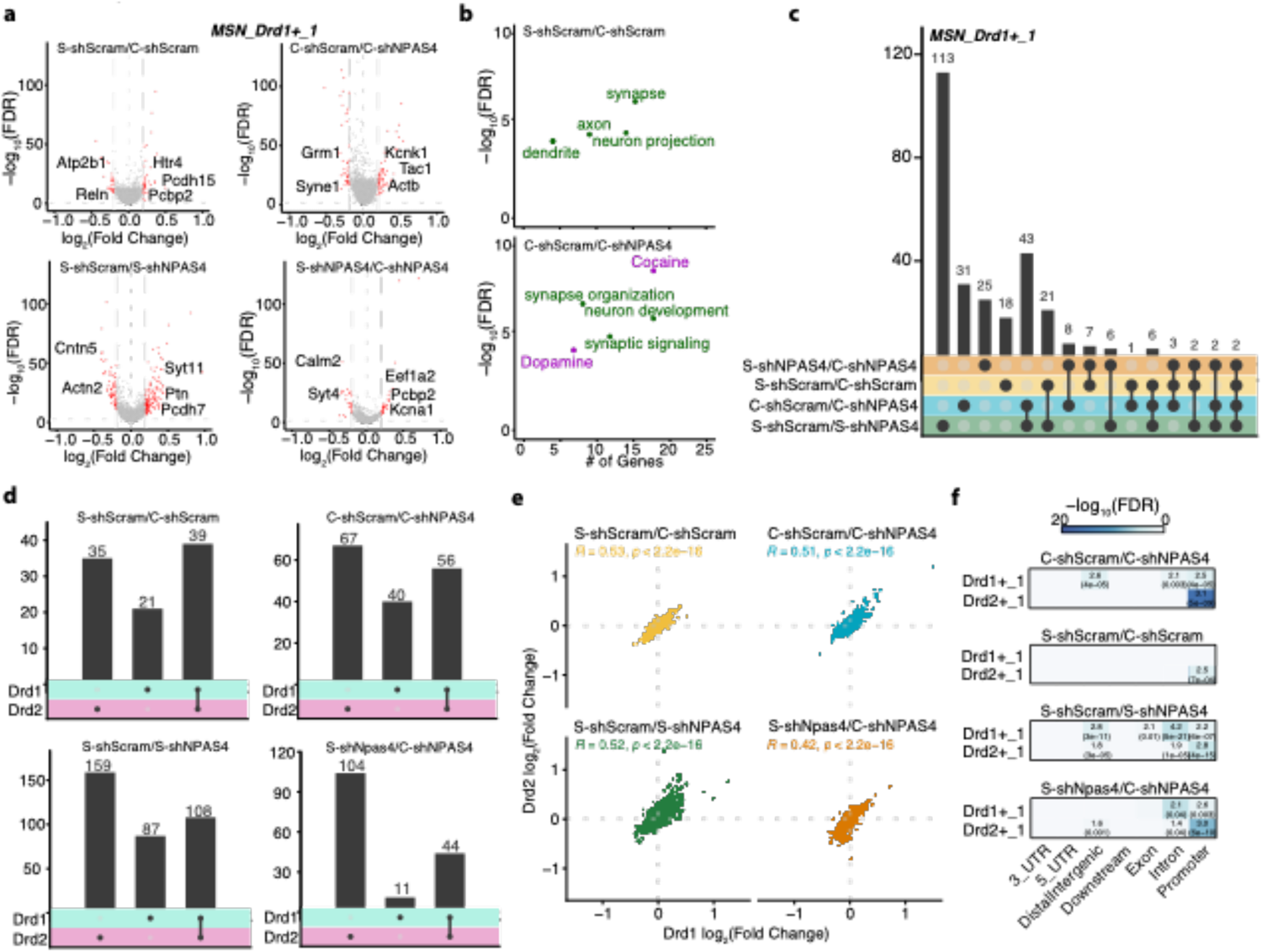
Single nuclei RNA-sequencing of the mouse NAc after NPAS4 knockdown and cocaine CPP: D1-MSN-specifc DEGs between group comparisons, overlapping DEGs in NAc D1-vs D2-MSNs, and comparison to published NPAS4 ChIP-seq data. **a**, Volcano plots depicting D1-MSN-specific DEGs in cocaine CPP vs saline CPP mice +/-NPAS4. **b**, Gene ontology analysis of DEGs in D2-MSNs comparing (top) “Saline CPP” vs “Cocaine CPP” and (bottom) “Cocaine CPP” vs “Cocaine CPP + shNPAS4.” X-axis depicts the number of genes in each category whereas the Y-axis correspond to the −log10(FDR) based on Fisher exact test. **c**, Upset plot showing the number of overlapping DEGs between groups in D1-MSNs. **d**, Upset plots showing DEGs specific to D1-MSNs, D2-MSNs, or both cell types. **e**, Correlation plots depicting the concordance between fold changes of DEGs in D1- and D2-MSNs and significant correlation in all 4 groups. Highlighted the correlation value and the relative p-value (Spearman rank correlation). **f**, Heatmaps depicting the enrichment of DEGs in D1- and D2-MSNs with NPAS4 ChIP-seq targets from Kim et al., 2010. Gradient color corresponds to the −log_10_(FDR). Heatmaps show the FDR value (parenthesis) and the odd ratios calculated with Fisher’s exact test.

**Extended Data Fig. 7:**
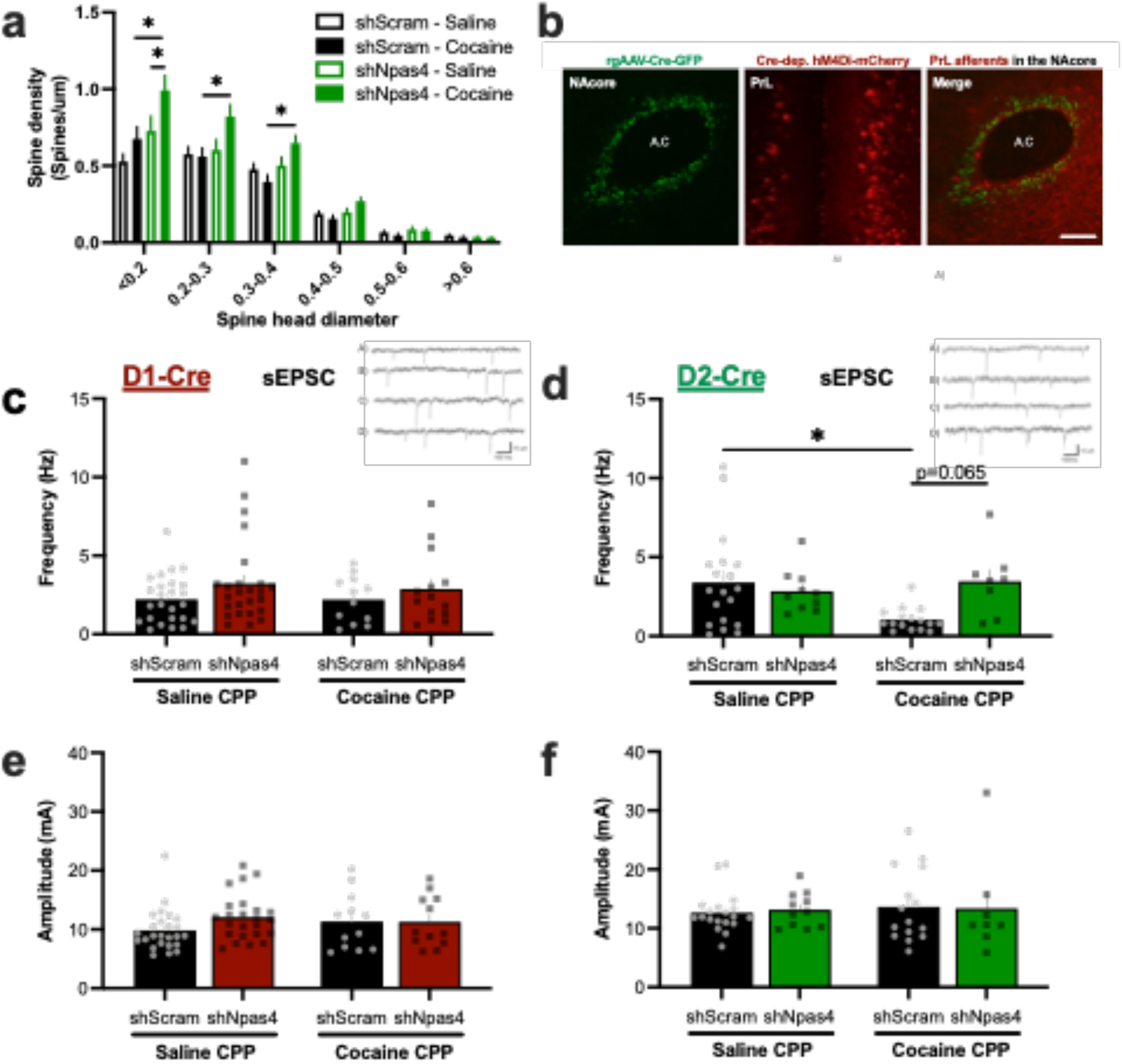
NPAS4 knockdown in D2-MSNs increases thin dendritic spines and modulates spontaneous EPSC frequency, but does not affect D1-MSN synaptic transmission. **a**, Quantification of spine density in D2-MSNs by spine head diameter. **b**, Representative image showing AAV-retrograde Cre in the nucleus accumbens core (green) and AAV-Cre-dependent hm4Di-mCherry in the prelimbic cortex (red). **c-d**, Spontaneous EPSC frequency in D1-MSNs and D2-MSNs. **e-f**, sEPSC amplitude after NPAS4 knockdown in D1-MSNs (left) and D2-MSNs (right) following cocaine conditioning. Insets on **c** and **d** show representative traces of spontaneous transmission in the following groups: (A) S-shScram, (B) S-shNPAS4, (C) C-shScram, and (D) C-shNPAS4. Data shown are mean ± SEM; *p < 0.05.

## Notes

### Competing Interest Statement

The authors have declared no competing interest.

