## Supplementary figures and images for "NPAS4 supports drug-cue associations and relapse-like behavior through regulation of the cell type-specific activation balance in the nucleus accumbens"

### Supplemental Figures

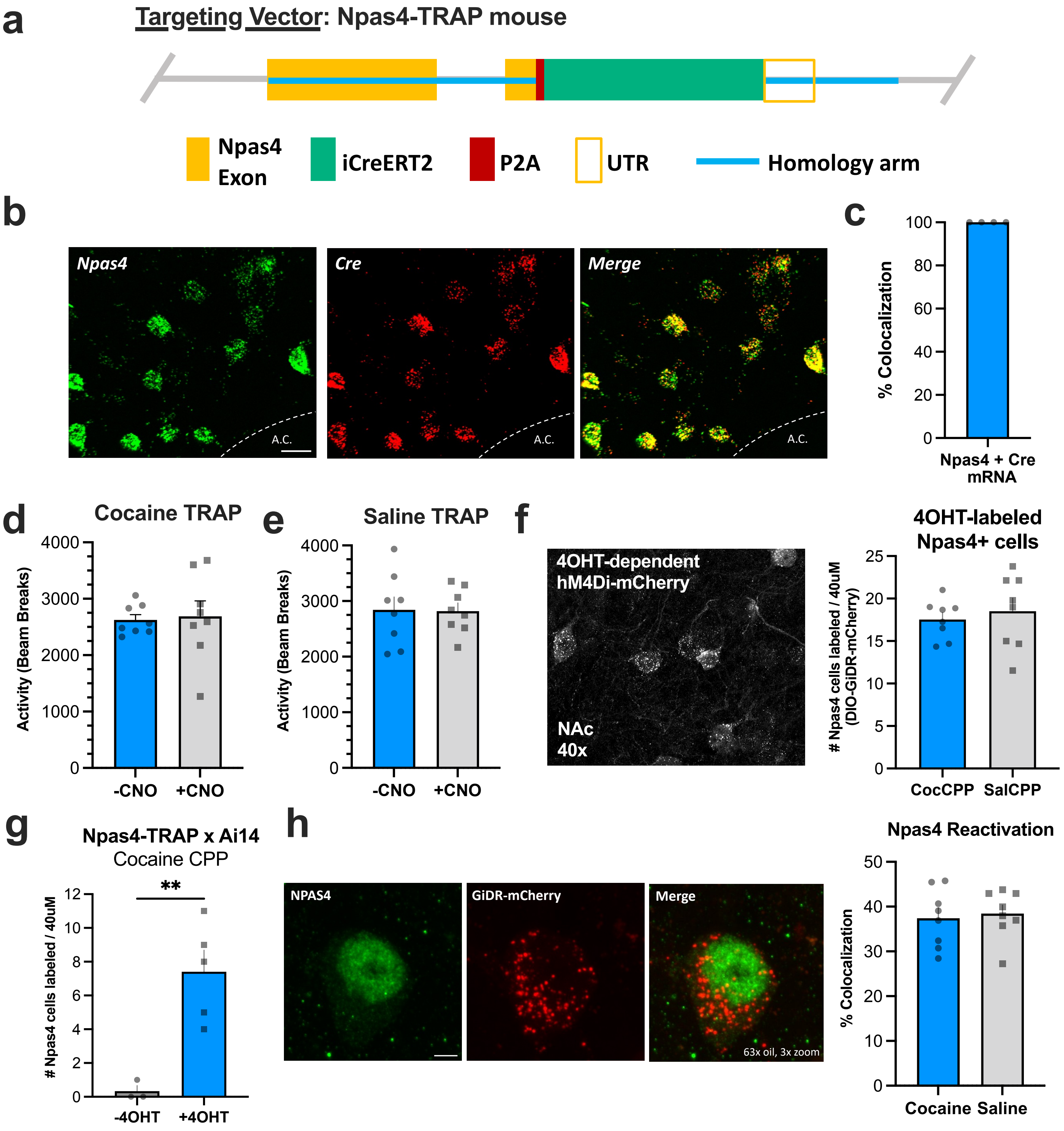

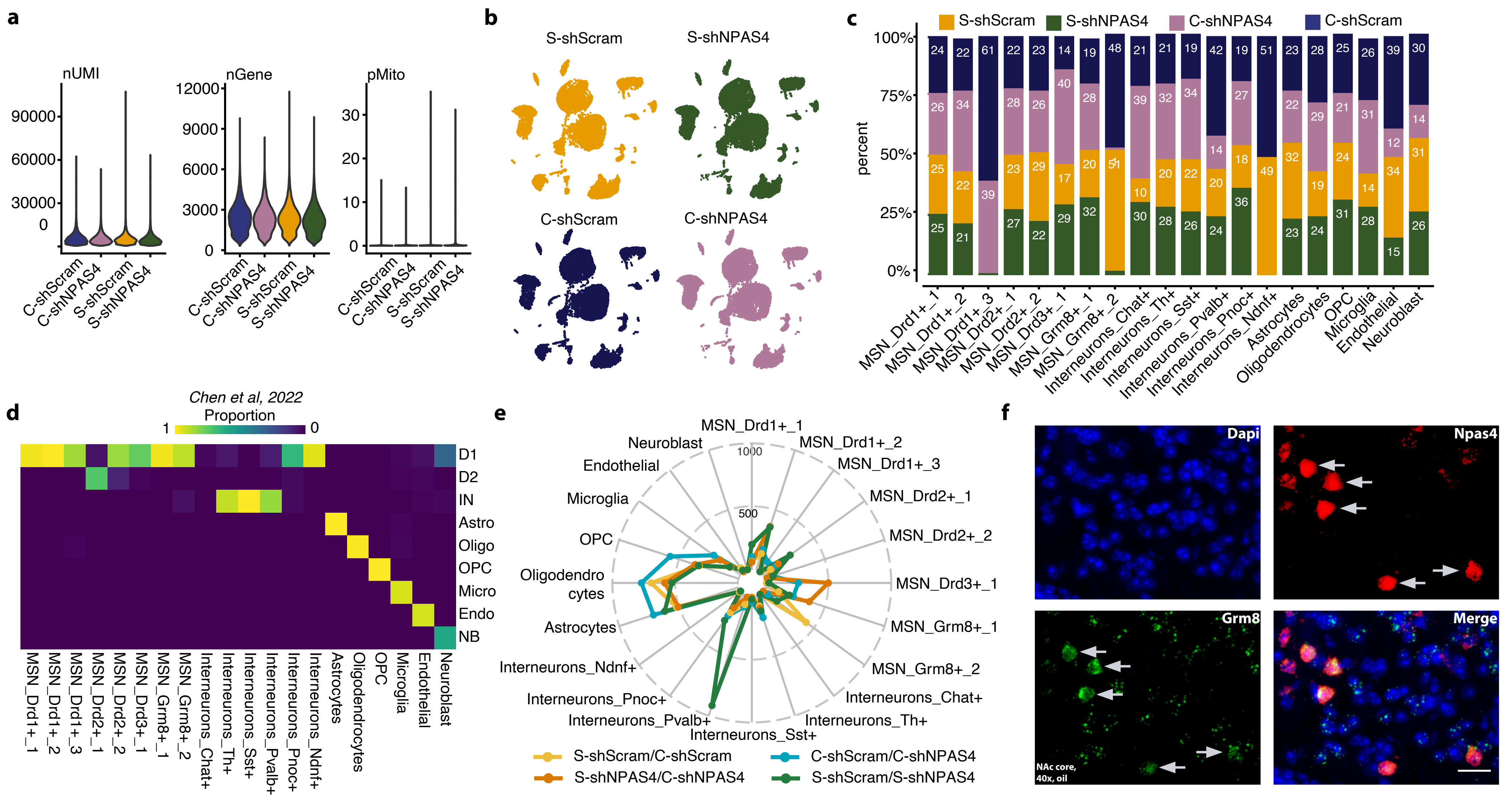

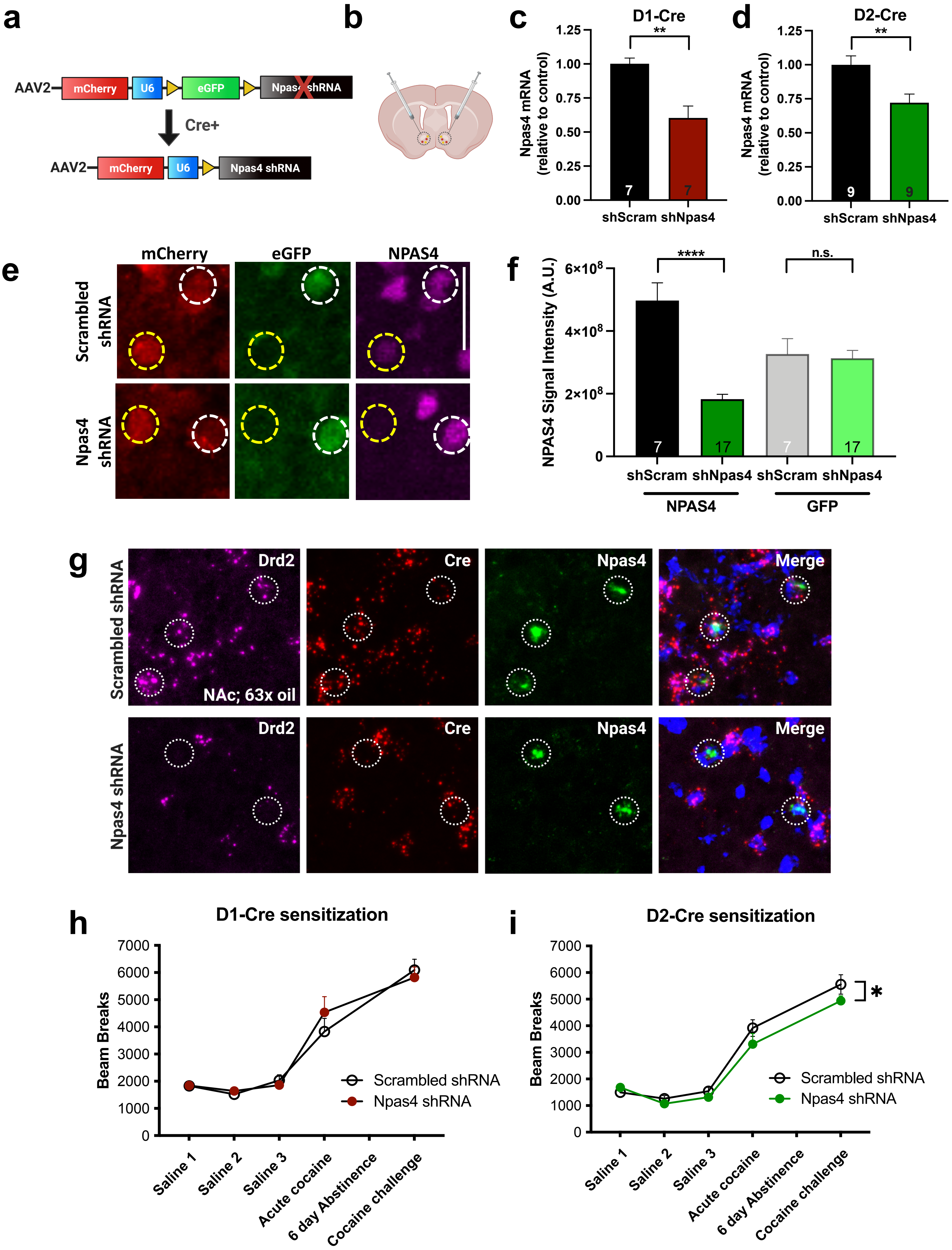

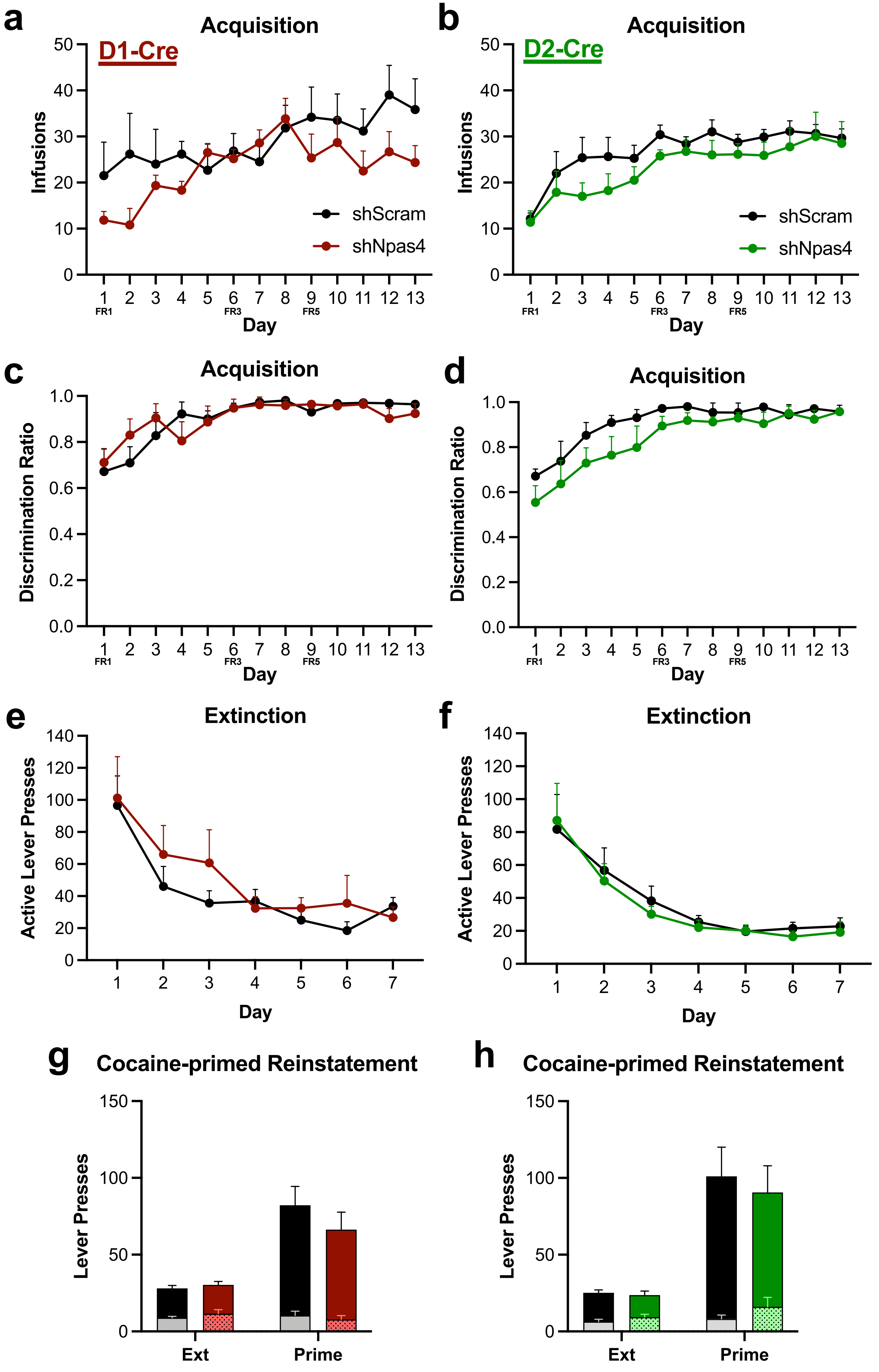

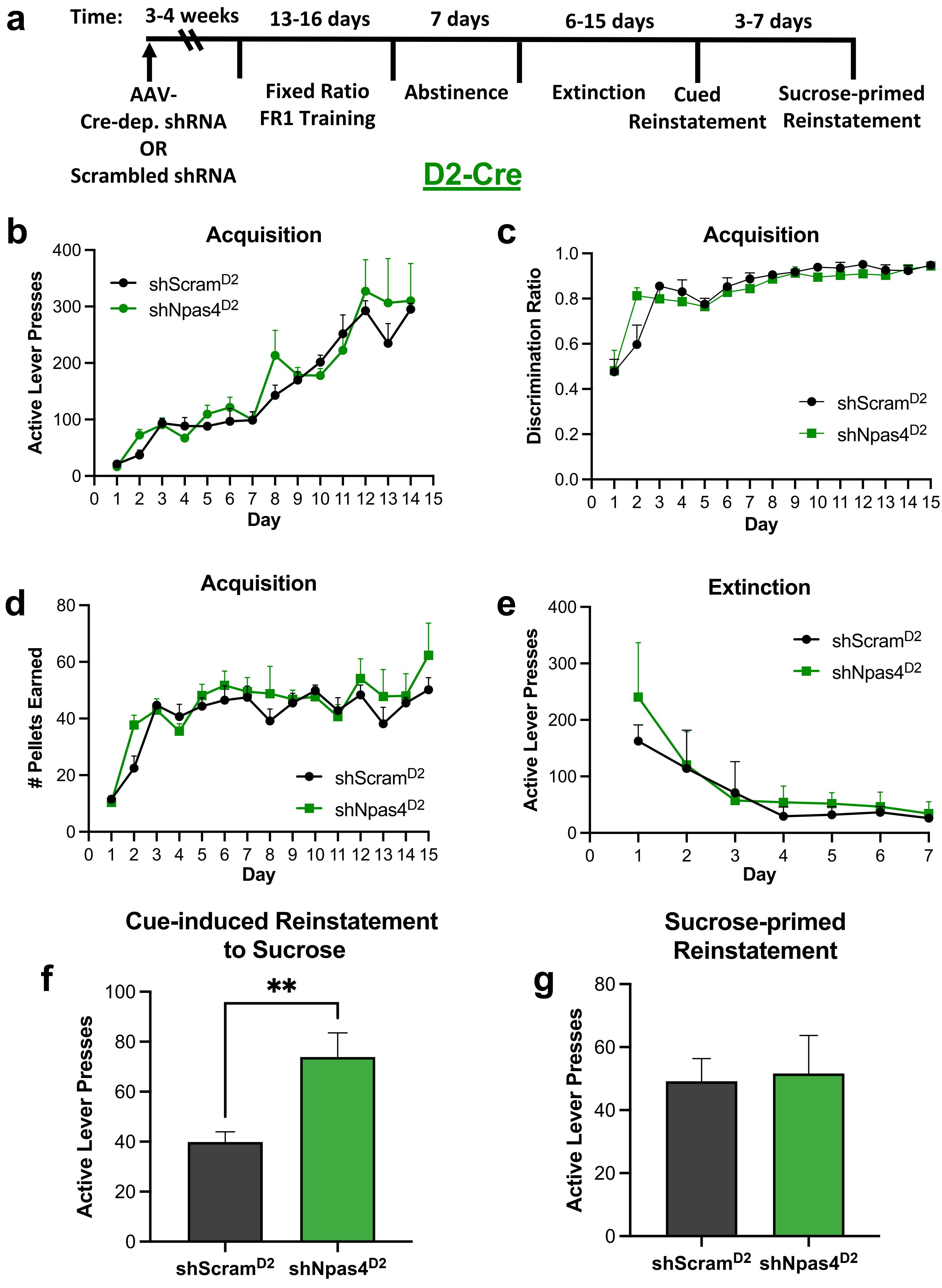

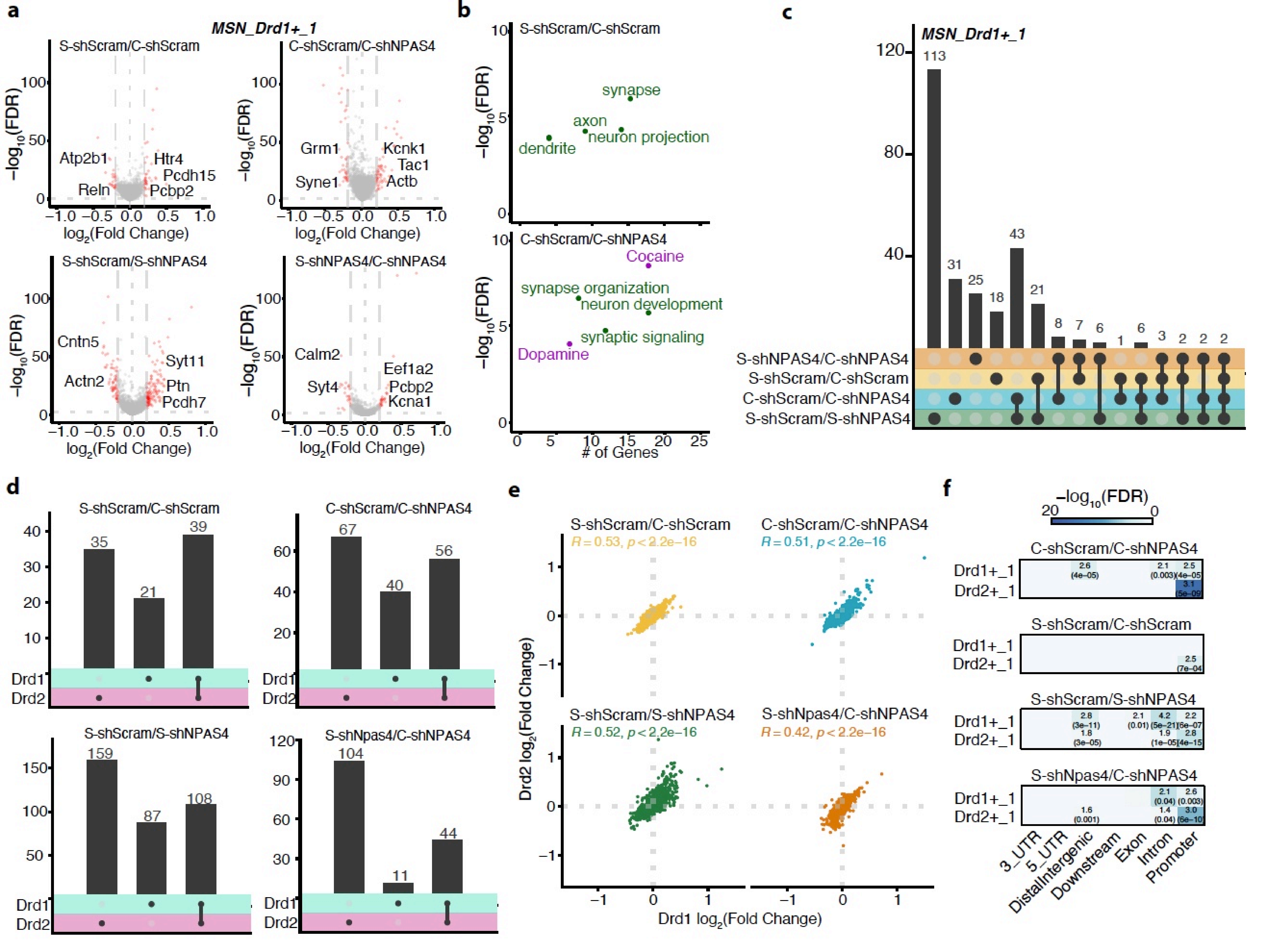

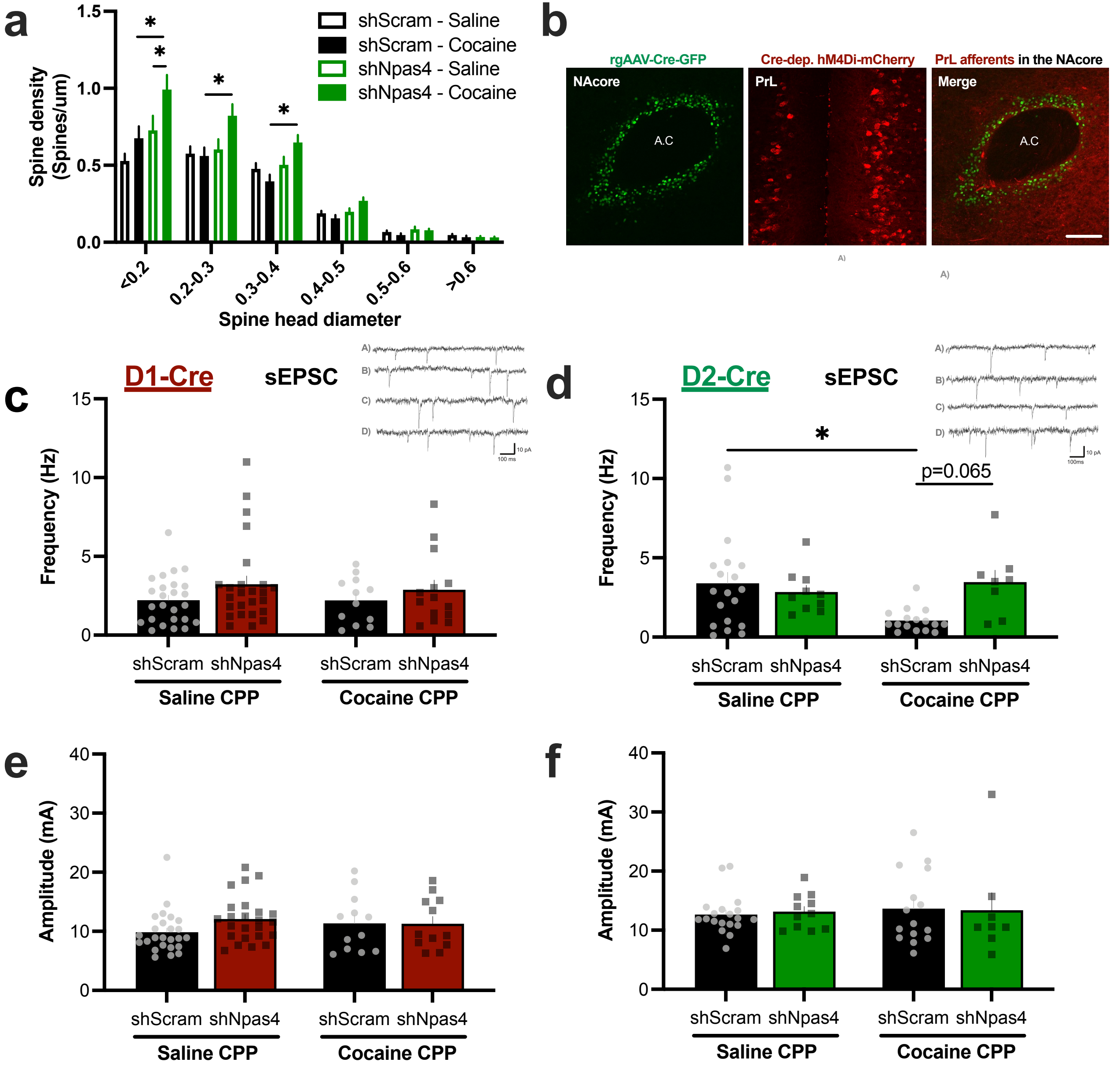
